# Fluid Uptake Pathways Are Differentially Modified in Rice (*Oryza sativa* L.) in Response to Drought and Salinity

**DOI:** 10.1101/2023.10.29.564576

**Authors:** Rukaya Amin Chowdery, H E Shashidhar, M K Mathew

**Author notes:** Correspondence: M K Mathew.

## Abstract

Salinity and drought adversely affect rice production globally. Here we have examined physiological responses to drought and salinity across four rice cultivars with varying sensitivity to these stresses. The salt tolerant Pokkali restricts fluid entry to limit Na^+^ uptake under saline stress, while the drought-tolerant ARB6 needs to enhance fluid uptake under drought. Surprisingly, Pokkali does reasonably well when subjected to drought as does ARB6 under saline stress - in contrast to the stress-sensitive but high yielding varieties IR-20 and Jaya. Both tolerant varieties use long roots to mine water under deficit conditions, increasing aerenchyma and suberization of the exodermis to provide oxygen to deep-reaching roots. Major alterations in patterns of suberization in both exodermis and endodermis are undertaken, the patterns being dramatically different under the two stresses. Genes implicated in suberin biosynthesis also showed variation in transcript levels under stress, corresponding with the observed suberization patterns. Osmolyte accumulation drives uptake of water under deficit conditions, while restricting fluid flow to symplastic routes minimizes Na^+^ entry. Overall, the morphological and physiological responses of the tolerant varieties ensure adequate fluid flow through the transpiration stream without excessive salt uptake, thereby promoting growth under both drought and salinity.

## Introduction

Rice (*Oryza sativa* L.), a dietary staple of more than half of the world, is generally cultivated in semi-aquatic conditions and is very salt sensitive [1–3]. Short term salinity leads to physiological drought and its persistence causes ionic stress [4,5] leading to altered K^+^/Na^+^ ratios, which affects all aspects of plant growth. Salinization of irrigated land leads to over a million hectares of land being lost to production annually. Salinization of arable land may have contributed to the demise of the early Mesopotamian civilization, making this a 6000 year old problem [6].

When grown in paddies with submersion, rice consumes an estimated 3000 to 5000 liters of water to produce a kilogram of grain [7]. Consequently, scientists have bred ―aerobic rice‖ [8–10], which does not require water logging [11]. Coarse, long roots, capable of branching and penetrating into the deep layers of soil have been selected for in breeding for drought tolerance [12–17]. Recent studies have shown aerobic rice to perform excellently in terms of high grain yield under severe drought stress [18,19]. However, the mechanism(s) employed to withstand drought have not been delineated. Nor is it clear as to whether traits that promote drought tolerance would adversely affect the ability of the plant to combat salt stress.

Some traditional varieties like Pokkali and Nona-Bokra are highly salt tolerant [20–22] whereas IR-20 is highly sensitive to salt [23]. Na^+^ compartmentalization in the shoot influences survival with high Na^+^ in the apoplast being correlated with poor survival [24–27]. Pokkali restricts Na^+^ loading into the xylem stream by building substantial apoplastic barriers in roots [28] and maintains low cytosolic Na^+^ [27,29] by reducing plasma membrane permeability to Na^+^ as well as sequestering Na^+^ in the vacuole [30]. It is not clear as to whether these mechanisms would be of use in combating drought stress.

The common apoplastic barriers in roots are casparian bands and suberin lamellae [31,32]. While Casparian Bands are deposited on radial and anticlinal walls of cells, suberin lamellae are deposited as secondary wall thickenings of the primary cell wall [32,33]. Elongases, hydroxylases and peroxidases are the important enzymes involved in suberin biosynthesis [34–36]. The role of suberin barriers in the response of higher plants to salinity, flooding, mechanical impedance, ion-deficiency and biotic stresses has been well studied [37–39]. Changes in suberin deposition following drought stress have not been investigated nor have the underlying molecular mechanisms been elucidated.

In the light of changing environmental conditions, salinity and drought are expected to create increasingly more severe challenges to rice production in the future [40]. Roots are directly and constantly in contact with soil and are responsible for water and nutrient uptake. Both anatomical and physiological strategies could foster growth and survival of the plant under stress. It would be of interest to investigate whether the mechanisms conferring tolerance to one abiotic stress (for example salinity) are beneficial or antagonistic to countering another abiotic stress such as drought. In the present study, we investigate the responses of four rice varieties to drought and salinity. We have chosen one specialist cultivar that has been shown to do well under drought – ARB6; another under salinity - Pokkali; one cultivar known to be very sensitive to both stresses – IR-20; and finally one that is moderately sensitive to both stresses - Jaya. All four were subjected to drought and salt in separate experiments. The expectation was that the anatomical and physiological responses would be distinct for each stress; that specialists would, in consequence, do well under ―their‖ respective stress, but would be sensitive to the other stress; and that the responses would be graded according to the degree of tolerance/sensitivity among the four cultivars studied. Given the range of sensitivity among the cultivars tested, such findings would set up hypotheses that could be subsequently tested.

## Materials and Methods

### Experimental setup and growth condition

Four cultivars of rice (Pokkali, ARB6, IR-20 and Jaya) were used based on their differences in tolerance and sensitivity to drought and salinity. Plants were grown and subjected to drought or salinity in PVC pipes of diameter 8cm and length 60cm. Field soil with pH of 5.7 was mixed with organic vermicompost (Organic carbon 9 – 17%, Nitrogen 0.5 – 1.5%, Phosphorus 0.1 – 0.3%, Potassium 0.15 – 0.56%, Sodium 0.06 – 0.3% and micronutrients) in the ratio 2:1 (soil: manure), and used to fill the pipes. Further compaction of the soil was achieved by watering and pressing the soil at intervals in order to mimic the compaction of soil in field conditions as described in [41]. Seeds were sown in three batches with 72 pipes in each batch. Plants were thinned to one per pipe and grown for 45 days with average temperature of 31.2 °C, relative humidity of 67.3%. In total, 216 pipes were used with 18 replications per variety and using 6 replicates for each analysis. Pipes arranged in a randomized complete block design were regularly watered till the 38^th^ day. From the 39^th^ day onwards, one of the batches was exposed to well-watered condition referred to as ―Control‖ and other two batches were exposed to either drought (no irrigation till the soil reached about 30-40% of field capacity) or salinity (150mM NaCl) stress for one week. Soil moisture content was monitored using moisture meter (Procheck, Decagon Devices Inc., USA) (Supplementary Information 1A, B, C, D). Electrical conductivity of soil under salinity condition was also measured (Supplementary Information 1E).

### Analysis of plant morphology and biomass

After one week of stress, the soil column along with the whole plant was taken out of the pipe. Roots were carefully washed free of soil. The number of lateral roots, tillers and leaves was counted manually and total shoot and root lengths measured. Fresh weight of the plants was recorded at this stage and dry weight after oven-drying at 70°C for 48 hours.

### Measurement of xylem sap exudation

Sap measurements were recorded using the method described [42,43] with some modifications. Rice varieties were grown in six replicates in PVC pipes and exposed to drought and salinity stress as described above. At 06:00 pm on the 45^th^ day, shoots were cut ∼5 cm above the soil surface. Cut stems attached to the root system were covered with pre-weighed blotting paper, then covered with polyethylene wrapper, which was in turn tightly sealed at the base with a rubber band (to avoid evaporation of the sap). The setup was left for 12 hours, followed by removing the blotting papers which were immediately weighed to quantify the amount of xylem sap sent up by the root system. The values were normalized to the root mass of the plant from which sap was collected. The blotting papers were then used to estimate the Na^+^ content of the sap.

### Measurement of root exudation in the absence (osmotic exudation) and in the presence of hydrostatic pressure gradients

Root *L*_pr_ measurements were carried out as described by [44], with some modifications. After excising the shoot at 5cm above the soil surface, the soil was carefully washed away from the roots and the root system submerged in a container of nutrient solution was placed in a pressure chamber. All tillers were closed using clamps except the main tiller which was threaded through a rubber stopper sealed with silicone sealant. Pre-weighed blotting paper was used to cover the cut surface of the main tiller to absorb the exuded xylem sap and was covered with a polyethylene wrapper to avoid evaporation. The blotting paper was weighed to determine the amount of xylem sap exuded by osmotic pressure. The osmotic pressure of the nutrient solution measured using a freezing point Osmometer (Osmomat 030, GONOTEC GmbH, Germany) was 0.0075 MPa. The measured reflection coefficient, σ_sr_ of the nutrient salts was 0.4. Plants used for measuring osmotic water flows through the root were also used to measure the hydrostatic water flow of the roots. The air in the chamber was pressurized and monitored with a pressure transducer. Pressure was slowly raised in steps of 0.05 MPa from 0MPa to 0.3MPa (±0.001 MPa). Sap flow was stable at pressures above 0.05MPa. At a given applied gas pressure (*P* in MPa), the volume exuded from the root system (*V* in m^3^) was plotted against time (Supplementary Information 2A). The slope of the *V* vs time curve, normalized to the surface area of the root system, yields the volume flow, *J*_vr_ in m s^−1^. Plots of *J*_vr_ against applied pressure were linear above 0.15 MPa. Root hydraulic conductivity (*L*_pr_) was estimated as the slope of the curve in this linear regime, illustrated with red line (Supplementary Information 2B).

### Estimation of crop canopy air temperature difference (CCATD)

Small plots were prepared in the field and seeds sown in five lines 15cm apart with ten replicates in each line. Control plots were watered regularly till 45^th^ day. Plants in the other set were well-watered for 38 days, following which the drought protocol was followed. At mid-day on the 45^th^ day, ambient and canopy temperature of both control and stressed plants was recorded by using AGRI-THERM III^TM^ infra-red sensing instrument (Everest Interscience Inc., USA). The difference in the two readings gives the Crop Canopy Air Temperature Difference (CCATD).

### Measurement of Photosynthetic rate

Photosynthetic rate was monitored using an infrared gas analyzer (IRGA) based LI-6400 photosynthesis system (LI-CoR Biosciences) on Day 0 (one day before subjecting the plants to drought and salinity stress) and during 7 days of stress period between 9.00 to 11.00 am, measurements were recorded on intact leaves 6 per plant using 6 plants per cultivar.

### Estimation of xylem sap Na^+^ content

Blotting papers with and without xylem sap were kept in deionized distilled water for one hour. The blotting paper was then discarded and the supernatant analyzed for Na^+^ ions using a flame photometer (Systronics Flame Photometer 128, Ahmedabad, India). The difference in the two values gives an estimate of Na^+^ in the xylem stream.

### Estimation of apoplastic Na^+^ and intracellular Na^+^ and K^+^ in shoot and root

The method described by [27] was used to release the total shoot Na^+^, apoplastic and intracellular fluid from shoot and root. The Na^+^ and K^+^ in the solution was estimated by flame photometry.

### Estimation of shoot and root osmotic adjustment

For the estimation of osmotic adjustment, shoot and root tissue was collected on the 45^th^ day. The tissue was immediately frozen in liquid nitrogen and brought to lab, where it was thawed and chopped into fine pieces with a razor blade. 500µL of deionized double distilled water was added to chopped tissue and centrifuged at 12000 rpm to collect the sap. The osmotic strength of the sap was estimated by using Osmometer (Osmomat 030, GONOTEC GmbH, Germany).

### Root microscopy

Thin sections (200µm) of Base (25 cm), Middle (15 cm) and Tip (5 cm) zones of roots were cut using a Mcllwain tissue chopper (The Mickle Laboratory Engineering Co. Ltd, UK). Sections were stained with 0.1% (w/v) berberin hemisulphate for one hour followed by 0.5% (w/v) aniline blue for 30 minutes as described by [45]. Stained sections were observed under Olympus FV1000 confocal microscope and imaged using 488nm excitation and 510-540nm emission. Suberin was seen as a fluorescent light yellow band shaped structure deposited around the endodermal and exodermal cells of the root, whereas Passage cells (PCs) showed no tangential suberin deposition. For counting and estimation of number of cells with or without suberin deposits, stained sections were observed using Nikon fluorescent microscope under UV light.

### Semi-quantitative RT-PCR analysis

Total RNA was isolated from roots of 45 day old plants using TRI-reagent (Sigma–Aldrich, St Louis, MO, USA) following the manufacturer’s instructions. RT-PCR was performed with 1 µg of RNA using 100 units of Moloney murine leukemia virus reverse transcriptase (Invitrogen) according to instruction of the manufacturer. The cDNA thus obtained was subjected to a 25-cycle PCR reaction with the following conditions: 3 min at 94°C followed by 25 cycles of 60 s at 94°C, 60 s at 58°C and 80 s at 70°C, and finally 5 min at 70°C. The constitutively expressed *Actin* was used as a control. The primer sequences and predicted amplicon sizes were 5’-CCTCTTCCAGCCTTCCTTCAT-3’(forward) and 5’- ACGGCGATAACAGCTCC TCT T-3’(reverse) for *Actin-1* (Os03g50890) (400 bp), 5’- CGCCTCACCTTCGATAACAT-3’ (forward) and 5’-CACTCGCAGTCCATTCTTCA-3’ (reverse) for *P450* (Os01g63540) (960 bp), 5’-TCGTAATCTTCTCCGCCATC-3’ (forward) and 5’- GATGTAGGCGAGCTCGTACC-3’ (reverse) for *Elongase* (Os03g12030) (821 bp), 5’- CCGTCTACCTCGTCGACTTC-3’ (forward) and 5’-ATCCATCCACGGGTTCTTCT-3’(reverse) for *Elongase* (Os02g11070) (1190 bp), 5’-TCGTCAATTGCAGCTTGTTC-3’ (forward) and 5’- TCTCCTTCGCTGGATTCACT-3’ (reverse) for *Elongase* (Os06g39750) (867 bp).

### Statistical analysis

The data in the figures have been presented as the mean values ± SE; n = 6. Student’s t-test was used to estimate the significant differences at P < 0.01 and P < 0.05, between control and stress treatments. For root hydraulic conductance experiment, analysis of variance (ANOVA) was used to detect the significant differences and Least Significant Difference (LSD) test was used to determine genotypic differences in each treatment and group them into letter classes.

## Results

We selected four cultivars of rice with differing sensitivities to drought and salinity for this study. All four varieties were grown in soil under well-watered conditions for 38 days, following which they were subjected to either drought or saline stress for a week. Plants were harvested on the 45^th^ day for analysis.

### Responses to drought stress

#### Morphophysiological changes

At the end of the one week drought regime, all of the experimental plants were alive. Soil moisture content, measured at a point 20 cm below the surface, declined for all varieties in a similar manner (Supplementary Information 1A, B, C, D). IR-20 and Jaya plants showed clear signs of distress: leaf wilting and chlorosis; whereas ARB6 plants appeared quite healthy and Pokkali plants were not severely affected (Figure 1A, B). Photosynthetic rates for all 4 varieties were the same on Day 38 – i.e, the day before stress was imposed. The rates did not change substantially over the next 7 days for plants that were well watered (Figure 1C). The plants subjected to drought, however, showed a decline in photosynthetic rate that was extreme for IR-20, large for Jaya and moderate for ARB6 and Pokkali. ARB6 plants were stimulated to grow under drought and exhibited a substantial increase in biomass compared to control plants (Supplementary Information 3A). Pokkali plants showed no significant growth arrest, while IR-20 and Jaya displayed significant decline in growth (Supplementary Information 3A) with final biomass about 30% lower than in control plants.

**Figure 1.**
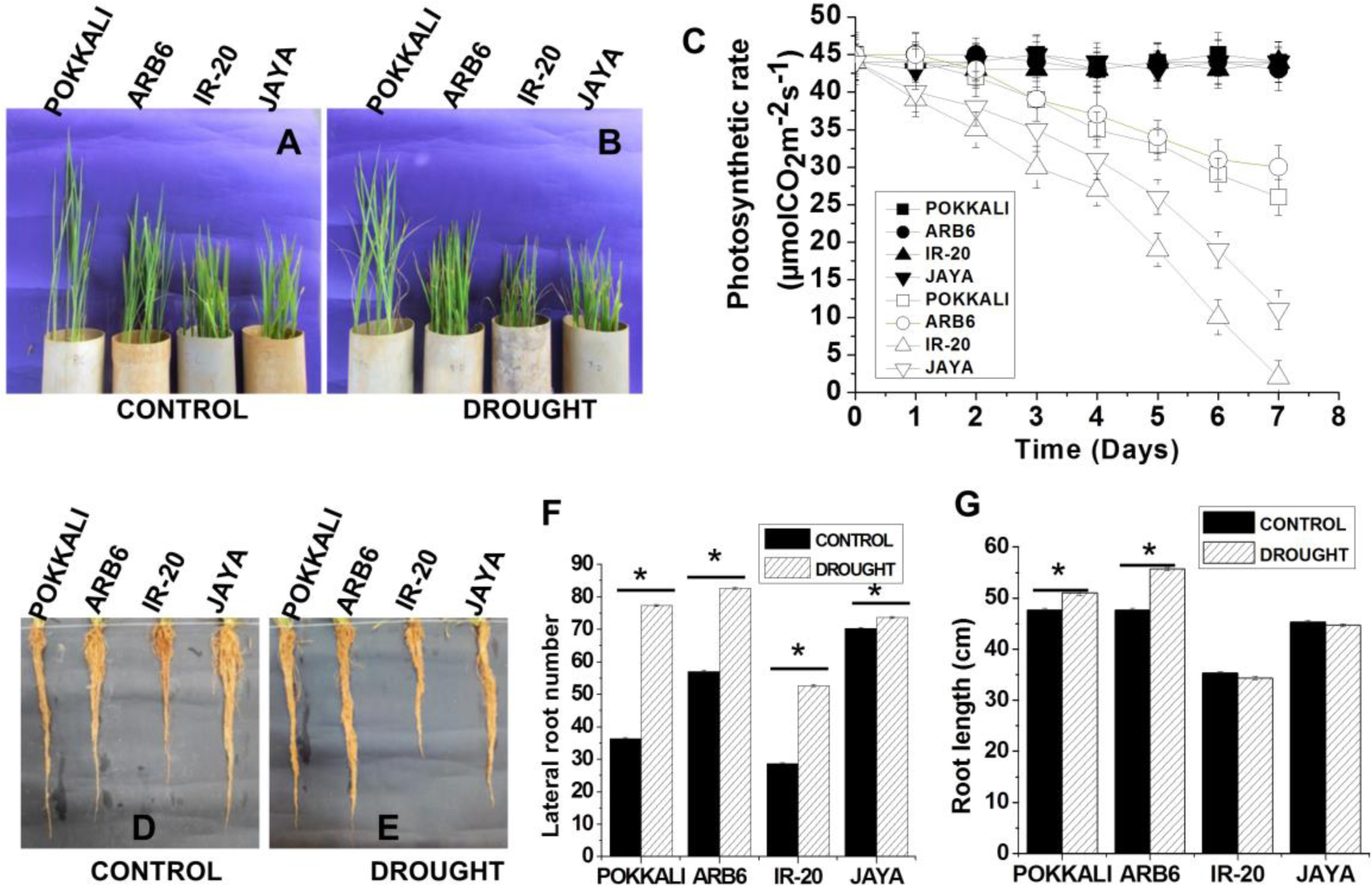
Rice varieties and their roots after being subjected to drought: Plants were grown in PVC pipes for 38 days and then stressed with drought (no irrigation) for a week. (A, D) Well-watered control and (B, E) drought stressed plants. (C) Photosynthetic rates were monitored throughout the stress period (Closed symbols - control and open symbols - drought) (F) Root number and (G) Root length. Data represents mean (+SE; n = 6), Asterisk indicates differences between control and drought with P < 0.05.

In the shoot, leaf number increased dramatically in ARB6 and marginally in Pokkali with declines observed in IR-20 and Jaya (Supplementary Information 3B). No significant change in plant height was displayed by any genotype (Supplementary Information 3C). Pokkali exhibited no change in tiller number, whereas the other three varieties exhibited increases under drought (Supplementary Information 3D). Unlike the shoots, roots of all four varieties appeared healthy with enhancement of root number (Figure 1D, E, F), the most prominent being Pokkali where the root number almost doubled. Root lengths showed modest increases (Figure 1G) of 20% in ARB6, 10% in Pokkali and none in IR-20 and Jaya. Aerenchyma were extensive in the basal region of control roots of all four varieties. However, under drought stress, these took over a large fraction of root area and extended to the middle and tip regions of ARB6 and Pokkali (Supplementary Information 4A). The data for the middle region of the root is presented in Figure 2A. Data for the base and tip regions are presented in Supplementary Information 4B, C.

**Figure 2.**
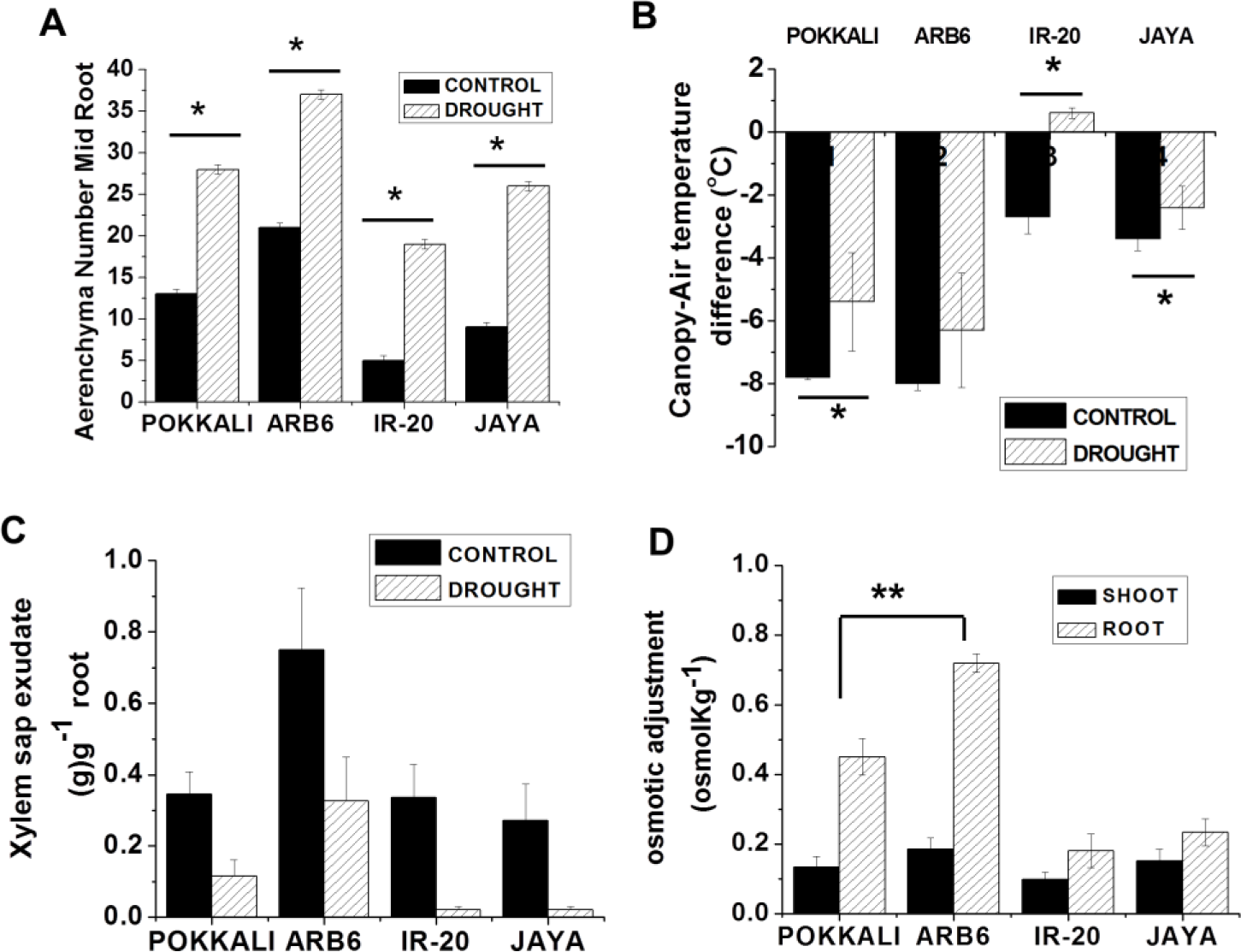
Plants were grown in PVC pipes for 38 days and subjected to control (well-watered), or drought (no irrigation) for a week. (A) Aerenchyma number of plants at the end of treatment period. (B) Crop canopy air temperature difference (CCATD). Ambient and canopy temperature was measured by using infra-red sensing instrument. (C) Xylem sap exudation. (D) Osmotic adjustment in shoot and root under drought condition. Data represents mean (±SE; n = 6), Asterisks indicate differences between control and stressed at *P < 0.05 and **P < 0.01.

Overall, the analysis of gross morphological features demonstrates that ARB6 plants do significantly better over a one week of drought stress protocol than control plants, whereas IR-20 coped poorly. Jaya exhibited moderate sensitivity to drought. Interestingly, Pokkali which was selected as a salt tolerant variety and reported to display characters that could sensitize it to drought, actually does well under this stress.

#### Canopy temperature

Increase in biomass can be achieved either by increasing water use efficiency or by accessing additional water sources under drought. Evaporative cooling of leaf tissue utilizes transpired water and reduces canopy temperature below that of the ambient air. The magnitude of this reduction in temperature is thus a good indicator of the extent of transpiration. Canopy temperatures were around 8°C below ambient in ARB6 and Pokkali under control conditions and about 4°C in Jaya and IR-20 (Figure 2B). Canopy temperature differences declined under drought by 1 to 3°C in ARB6, Pokkali and Jaya, though the decline in ARB6 was not significant. The canopy of IR-20 was at ambient temperature under drought indicating that transpirational cooling was no longer active. Maintenance of large canopy temperature differences in ARB6 and Pokkali is indicative of high transpiration rates, suggesting that the plants were able to access adequate water sources even under drought conditions. Jaya maintained a small but significant canopy temperature difference under drought.

#### Xylem sap exudation

Measurement of the rate of xylem sap exudation confirms the hypothesis that ARB6 and Pokkali were able to access water under the experimental drought conditions. Exudation rates for Pokkali, Jaya and IR-20 were comparable under control conditions (Figure 2C). ARB6 had almost double this rate of sap exudation under control conditions. Drought dramatically reduced exudation rates in all four varieties, but the final exudation rate in ARB6 was comparable to that exhibited by the other three varieties under well-watered conditions (Figure 2C). Pokkali sap exudation was reduced considerably from well-watered conditions but is still significant. Very little sap was exuded by IR-20 and Jaya using root pressure alone.

#### Osmotic adjustment

Osmolyte accumulation under stress was estimated in terms of osmolarity differences with control sap. Very low osmotic adjustment was seen in the shoots of all varieties (Figure 2D). Both Pokkali and ARB6 exhibited large osmotic adjustment in the roots with the latter being significantly larger. Jaya exhibited moderate osmotic adjustment, while that of IR-20 was least.

#### Suberin deposition

Root hydraulics would be critical for delivering fluid to the growing shoot. We examined hydrophobic barriers on the endodermis and exodermis in all four varieties (Figure 3, Figure 4 and Supplementary Information 5). The percentage of fully suberized cells in the exodermis appeared comparable across all four varieties in control conditions (Supplementary Information 6A). IR-20 had a larger proportion of unsuberized cells than the other three varieties in all three regions analyzed – base, middle (mid) and tip of the root. After a week of drought, there was a significant decrease in the percentage of fully suberized cells in ARB6 and Pokkali with a concomitant increase in passage cells (PCs) (Supplementary Information 6B). The shift was most dramatic in ARB6. Taking the middle region of exodermis for example, completely suberized cells dropped from ∼30% to 20% while passage cells increase from ∼3% to 15% with little change in unsuberized cells (Supplementary Information 6B). A similar trend of decrease in suberized cells was observed in the base and tip regions as well, with a corresponding increase in passage cells (Supplementary Information 6B). No such decrease in completely suberized cells was seen in Jaya or IR-20 (Supplementary Information 6B).

**Figure 3.**
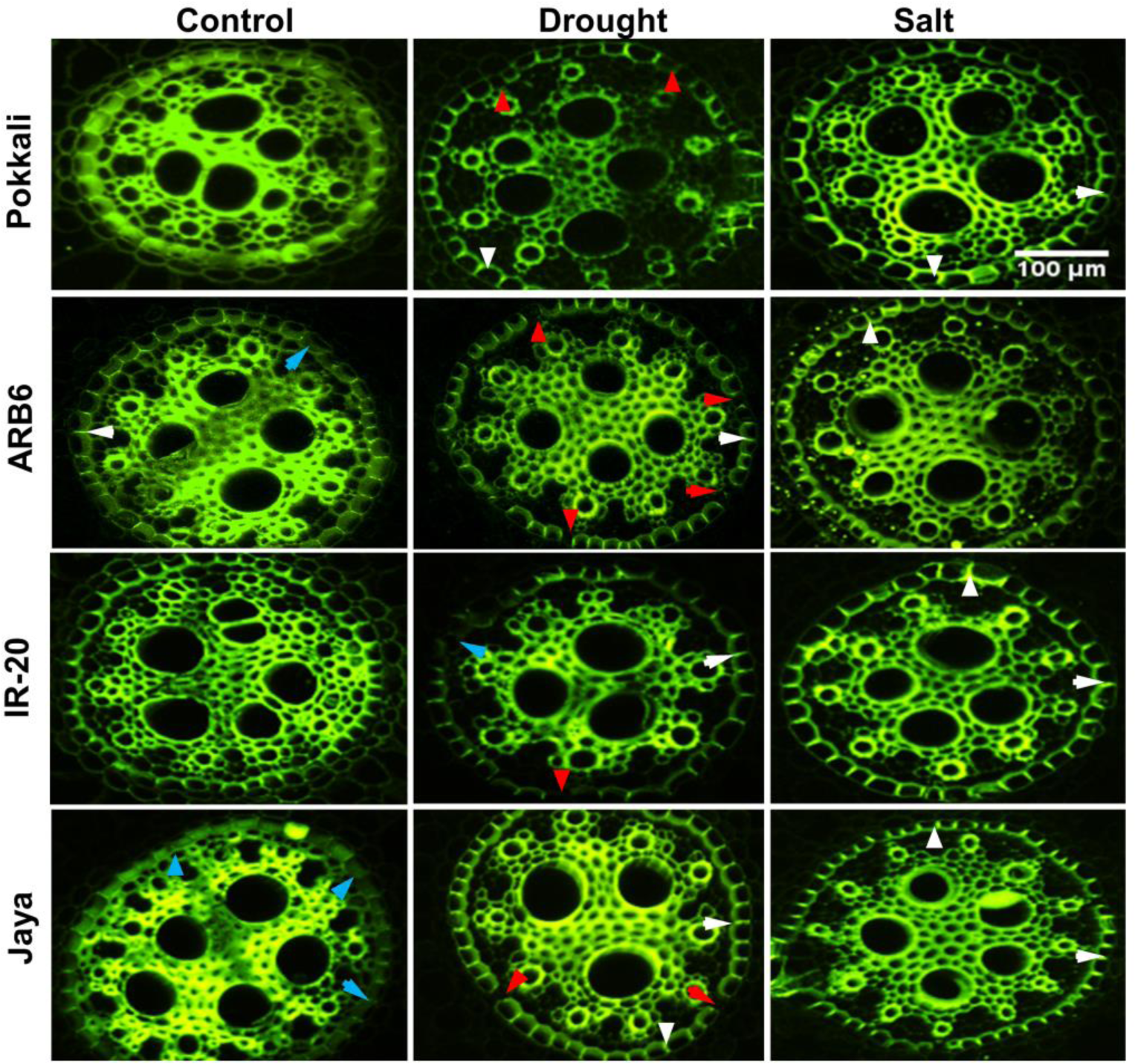
Innages of root sections showing unsuberized cells, passage cells and suberin deposits in root endodermis of plants grown in PVC pipes for 38 days and subjected to control (well-watered), drought (no irrigation) or salt (150 mMNaCI) for a week. Roots were washed, divided into three zones (Tip - 5cm, Mid - 15cm and Base - 25cm) from the root tip and cut into thin sections (200pm), then stained with berberine-aniline blue and imaged using Olympus FV1000 confocal microscope, 488nm laser was used for excitation. Red, white and cyan arrows show passage cells (cells with no tangential suberin deposition), suberin deposits and unsuberized cells respectively. Scale bar 100pm

**Figure 4.**
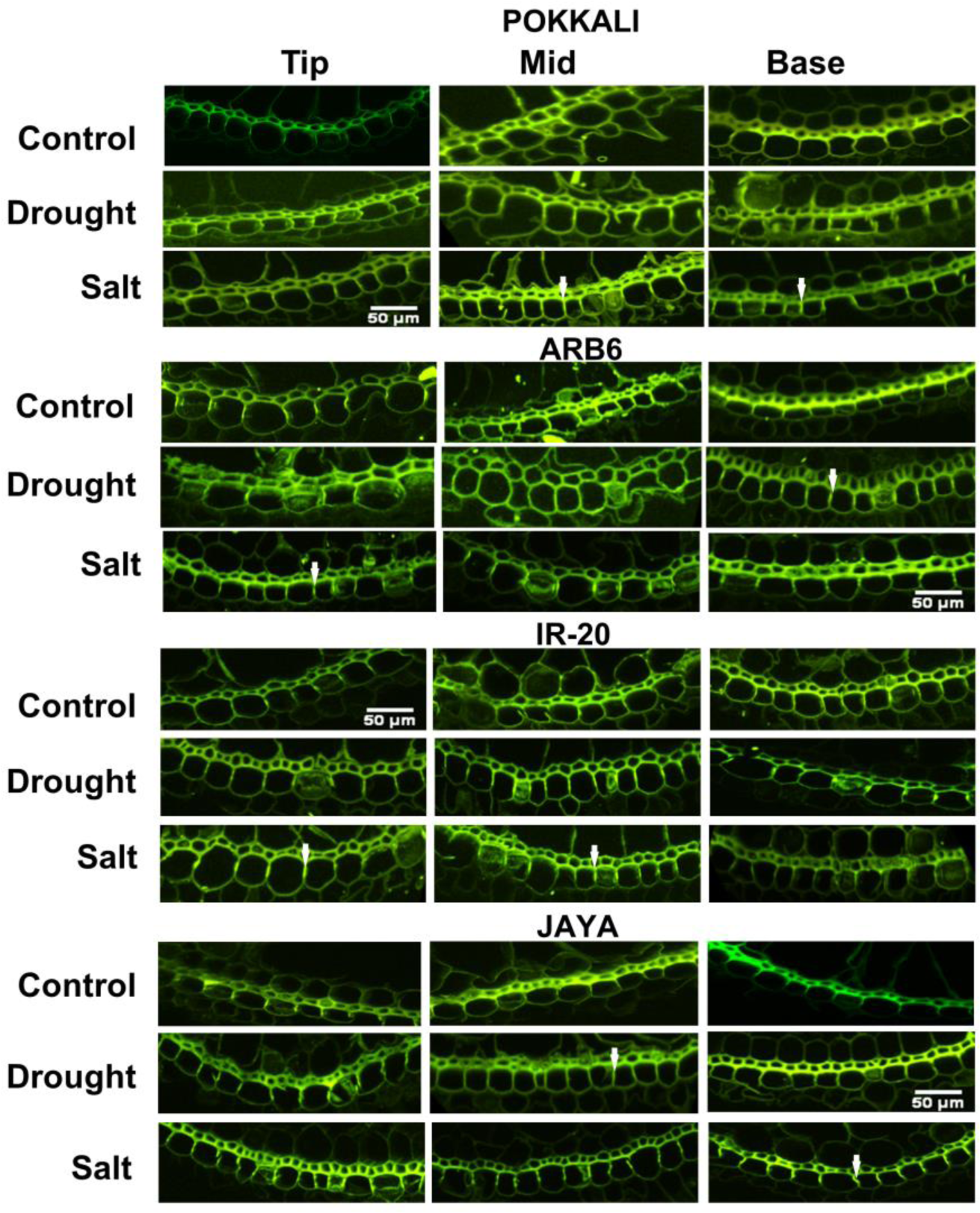
Images of root sections showing unsuberized cells, passage cells and suberin deposits in root exododermis of plants grown in PVC pipes for 38 days and subjected to control (well-watered), drought (no irrigation) or salt (150 mMNaCI) for a week. Roots were sectioned, stained and imaged as in Figure 3. White arrows show suberin deposits. Scale bars 50pm

The change in the pattern of suberization follows the same trend in the endodermis, only more so. Following drought, there was a significant decline in the percentage of completely suberized cells in both Pokkali and ARB6 but not in IR-20 and Jaya (Figure 5A, D, G). On the other hand, the percentage of passage cells in the Pokkali and ARB6 endodermis was comparable but greater than IR-20 and Jaya (Figure 5B, E, H). The changes in IR-20 and Jaya were relatively small and statistically insignificant in most cases (Figure 5B, E, H). No change was seen in percentage of unsuberized cells in Pokkali, ARB6 and Jaya. However, a decreasing trend was seen in IR-20 (Figure 5C, F, I). The tolerant plants thus increase the number of cells potentially available for uptake of fluid into the symplastic stream under drought stress.

**Figure 5.**
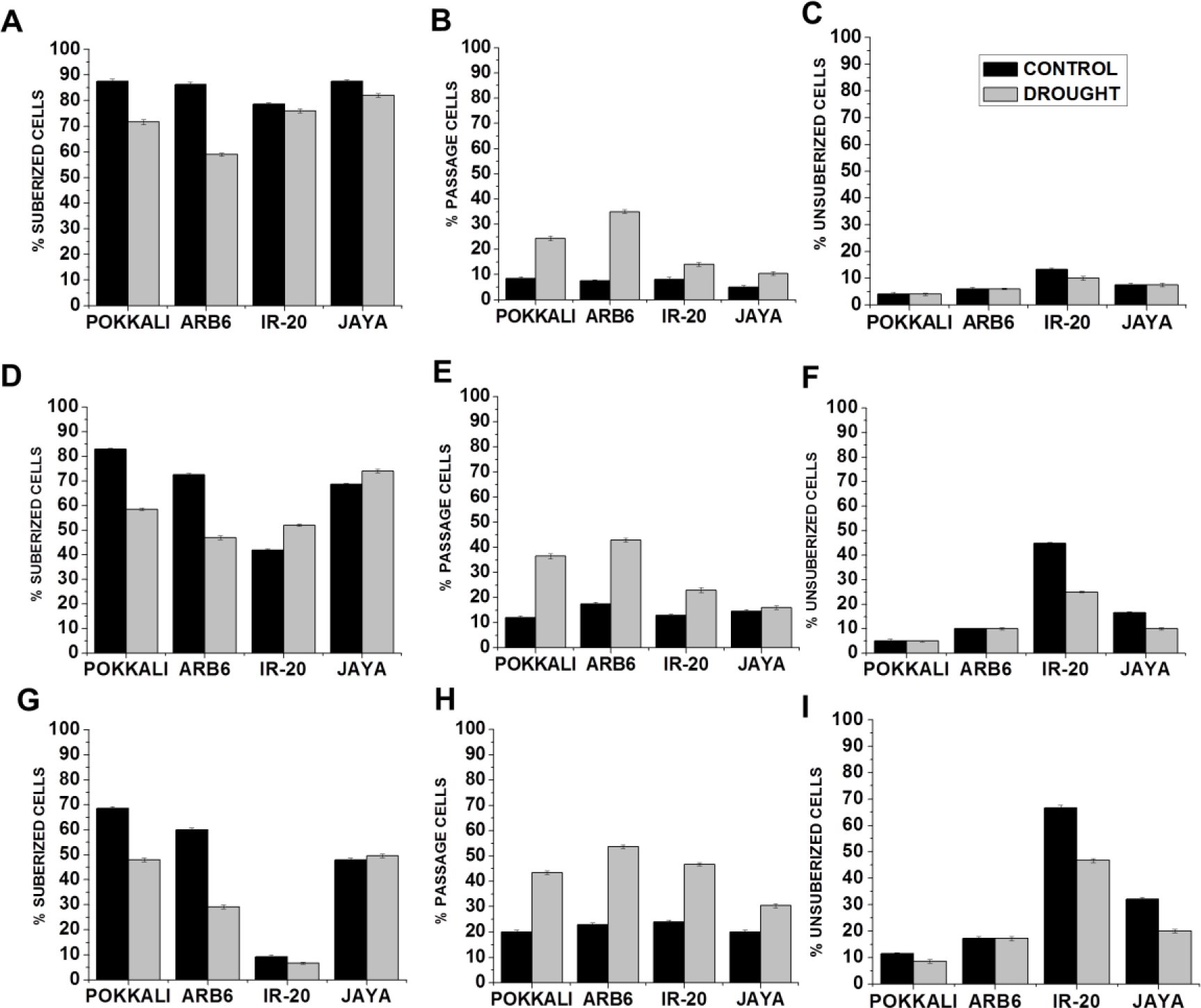
Quantification of suberization patterns in the endodermis to show suberized cells (having suberization on both sides and tangential areas), passage cells (with no tangential suberization) and unsuberized cells (without any suberin deposition) at (A - C) Base (D - F) Middle and (G - I) Tip of rice roots grown in control (well-watered) and drought (no irrigation) condition. Data represents mean (±SE; n = 6)

#### Root hydraulic conductivity (*L*_pr_)

Osmotic conductivity (*L_0_*) of Pokkali and ARB6 exceeded that of Jaya and IR-20 under control conditions, with little difference between the latter two (Table 1). While there was a trend towards increasing conductivity in all four varieties under drought, the changes were not significant in any case.

**Table 1.** Root hydraulic conductivity driven by external hydrostatic pressure (*L*_pr_) or by osmotic pressure alone (*L*_0_). For *L*_pr_, Pneumatic pressures were applied to the root medium to vary the driving force, while, no pressure was applied during *L*_0_ measurements (mean ±SE, n = 6). Values within a column followed by the different letters are significantly different and the values followed by the same letter are not significantly different at P<0.05 by LSD test.

|  | Variety | Control | Drought | Salinity |
| --- | --- | --- | --- | --- |
| Hydrostatic | Pokkali | 41±1.11c | 47±1.03c | 18±1.06a |
| $L_{pr}$ ($10^{-8}$ m MPa$^{-1}$ s$^{-1}$) | ARB6 | 49±1.16d | 56±1.05d | 23±1.08b |
| | IR-20 | 31±1.05a | 34±1.14a | 27±1.11c |
| | Jaya | 37±1.08b | 42±1.12b | 22±1.1b |
| Osmotic | Pokkali | 5.2±0.1b | 5.6±0.3c | 1.7±0.4a |
| $L_0$ (ms$^{-1}$) | ARB6 | 5.4±0.6b | 6.0±0.2c | 2.2±0.3a |
| | IR-20 | 4.1±0.1a | 4.3±0.3a | 2.8±0.5a |
| | Jaya | 4.4±0.3a | 5.1±0.1b | 2.2±0.4a |

Under control conditions, hydrostatic conductivity or *L*_pr_ was significantly higher in ARB6 than in the others, with Pokkali *L*_pr_ being greater than Jaya, which exceeded IR-20 (Table 1). Drought stress increased *L*_pr_ by 10 – 15% in the first three varieties, while the change in IR-20 was insignificant.

### Responses to salt stress

#### Morphophysiological analysis

At the end of a week of saline stress, all the experimental plants were alive. Pokkali plants looked healthy whereas all the other varieties appeared highly chlorotic (Figure 6A, B). Electrical conductivity of treated samples, measured at various depths, was significantly higher than in control pipes but similar across varieties (Supplementary Information 1E). Photosynthetic rates were the same for all four varieties on Day 38, and did not change over the next 7 days for plants that were well watered (Figure 6C). Rates fell sharply in all varieties when subjected to saline stress (Figure 6C). Both IR-20 and Jaya had very low photosynthetic rates on the 7^th^ days of stress, when the plants were harvested for further experiments. ARB6 decline was moderate while Pokkali decline was modest (Figure 6C).

**Figure 6.**
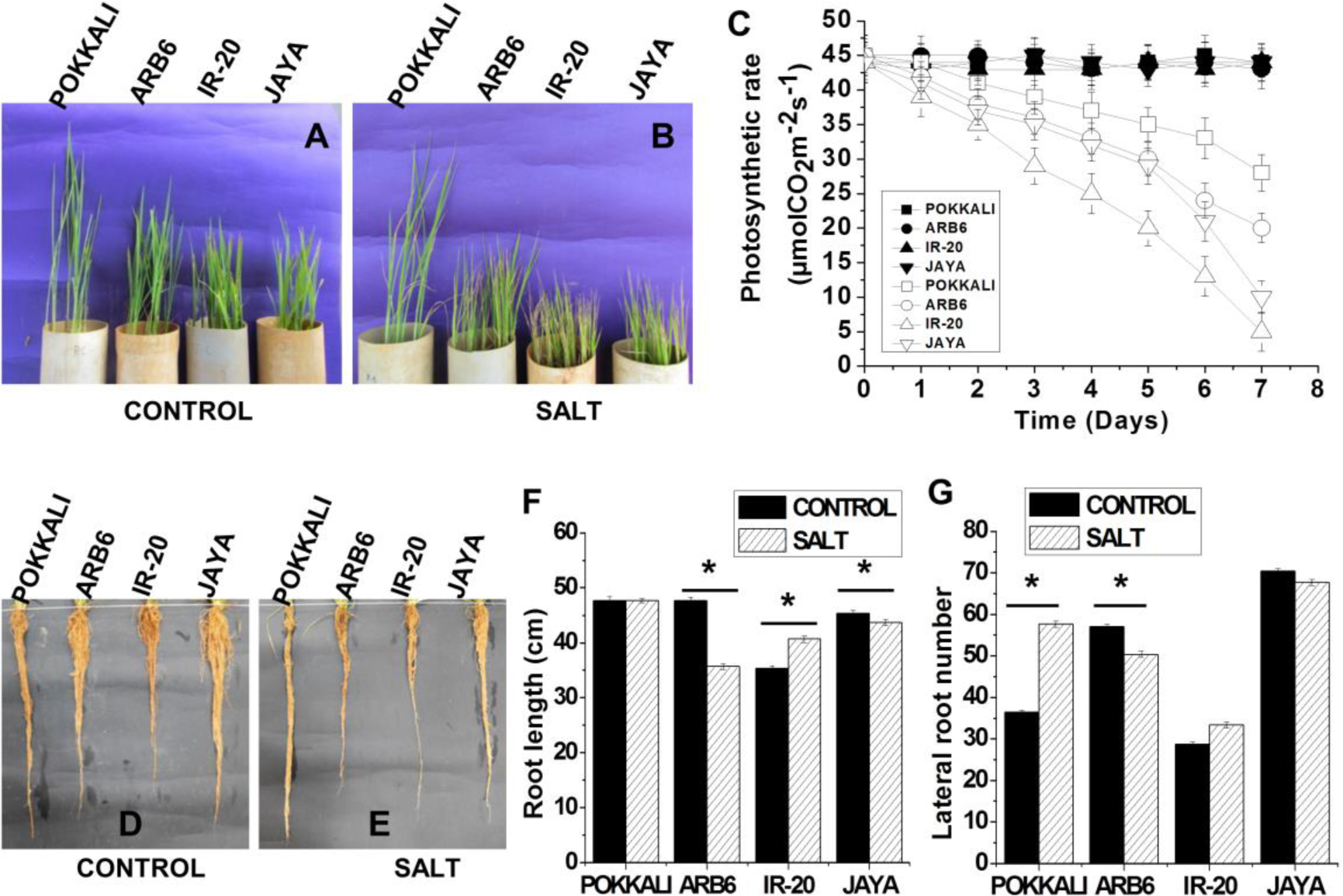
Rice varieties and their roots after being subjected to salt. Plants were grown in PVC pipes for 38 days and then stressed with salt (150mM) for a week. (A, D) control (B, E) Salt stressed plants. (C) Photosynthetic rate was monitored throughout the stress period (Closed symbols - control and open symbols - salt) (F) Root length (G) Root number of plants harvested on the 45th day. Data represents mean (±SE; n = 6), Asterisk indicates differences between control and salt that have P < 0.05.

The biomass of all varieties showed a significant decline of 30 to 45% (Supplementary Information 7A). No change in root length was seen in Pokkali as against significant declines in ARB6 and Jaya (Figure 6D, E, F). Surprisingly, IR-20 showed a significant increase in root length. The number of roots showed a significant increase in Pokkali plants, while they declined in ARB6 (Figure 6G). No significant change was seen in root number of Jaya and IR-20 (Figure 6G). No significant change in leaf number was seen in any variety (Supplementary Information 7B). Small but significant decreases in plant height were observed in Pokkali and ARB6, whereas no change was observed in Jaya and IR-20 (Supplementary Information 7C). Despite the decrease in plant height observed in both Pokkali and ARB6, tiller number increased in both varieties (Supplementary Information 7D). No change in tiller number was seen in Jaya or IR-20 (Supplementary Information 7D). Aerenchyma increased in all regions of all four varieties under saline stress, except in the tips of IR-20 roots (Supplementary Information 4D). Aerenchyma in the middle region of root is presented in Figure 7A. Overall, the morphological analysis confirms that only Pokkali performed well under salinity conditions. Interestingly, ARB6, which was selected as a drought tolerant variety, performed less poorly than Jaya, which is reported to be moderately tolerant to salt [27]. IR-20 did very poorly under salt stress.

**Figure 7.**
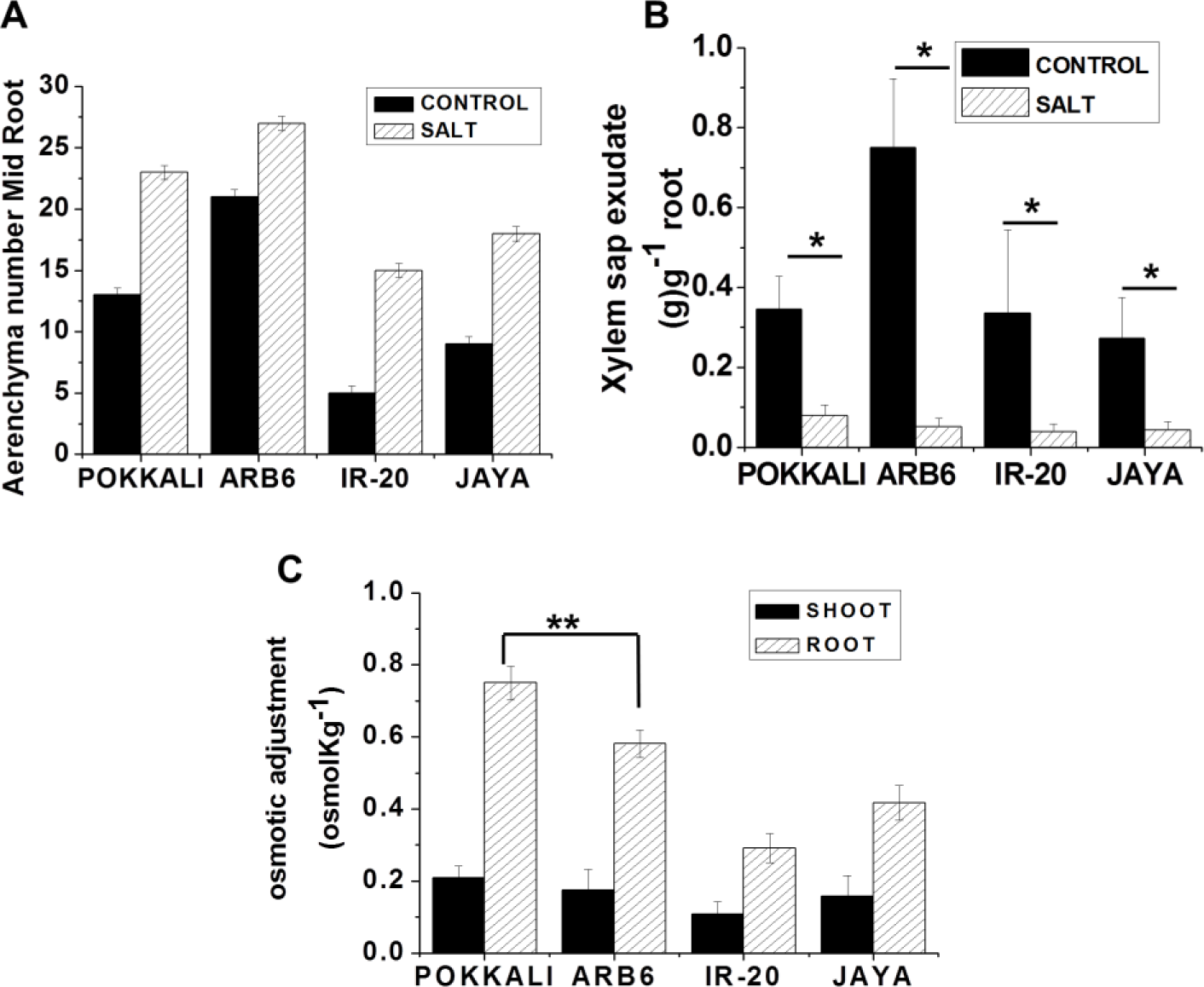
Plants were grown in PVC pipes for 38 days and then subjected to control (well-watered) or salt (150mM) for a week. (A) Aerenchyma number (B) Xylem sap exudate. (C) Osmotic adjustment in shoot and root under salinity condition. Data represents mean (±SE; n = 6), Asterisks indicate differences between control and stressed at *P < 0.05 and **P < 0.01.

#### Xylem sap exudation and osmotic adjustment

The extent of xylem sap exudation reduced dramatically in all four varieties, with only Pokkali showing significant exudation following saline stress (Figure 7B). Very low osmolyte accumulation was seen in the shoots of all varieties (Figure 7C). Osmolyte accumulation was significant in the roots of all the varieties with Pokkali accumulating significantly more osmolytes than ARB6, while moderate adjustments were seen in Jaya and IR-20 (Figure 7C).

#### Suberin deposition

The extent of suberization increased in all four varieties and in all regions of both exodermis (Supplementary Information 6C) and endodermis (Figure 8A, D, G). The changes were most dramatic in the endodermis of Pokkali. A few endodermal cells lacked suberin deposits in their tangential regions. These cells were adjacent to xylem elements and have been taken to be passage cells. An increase in passage cells was seen in all varieties except Pokkali where there was a decrease (Figure 8B, E, H). No unsuberized cells were seen in the endodermis in either the base or mid regions of any variety except IR-20 (Figure 8C, F, I), whereas unsuberized cells were observed in the tip region in all four varieties, where over 30% of cells in IR-20 are unsuberized (Figure 8C, F, I). It may be noted that even in the tip region, Pokkali and ARB6 had very few unsuberized cells and Pokkali had few passage cells in this region. The other three varieties had comparable fraction of passage cells in the tip region.

**Figure 8.**
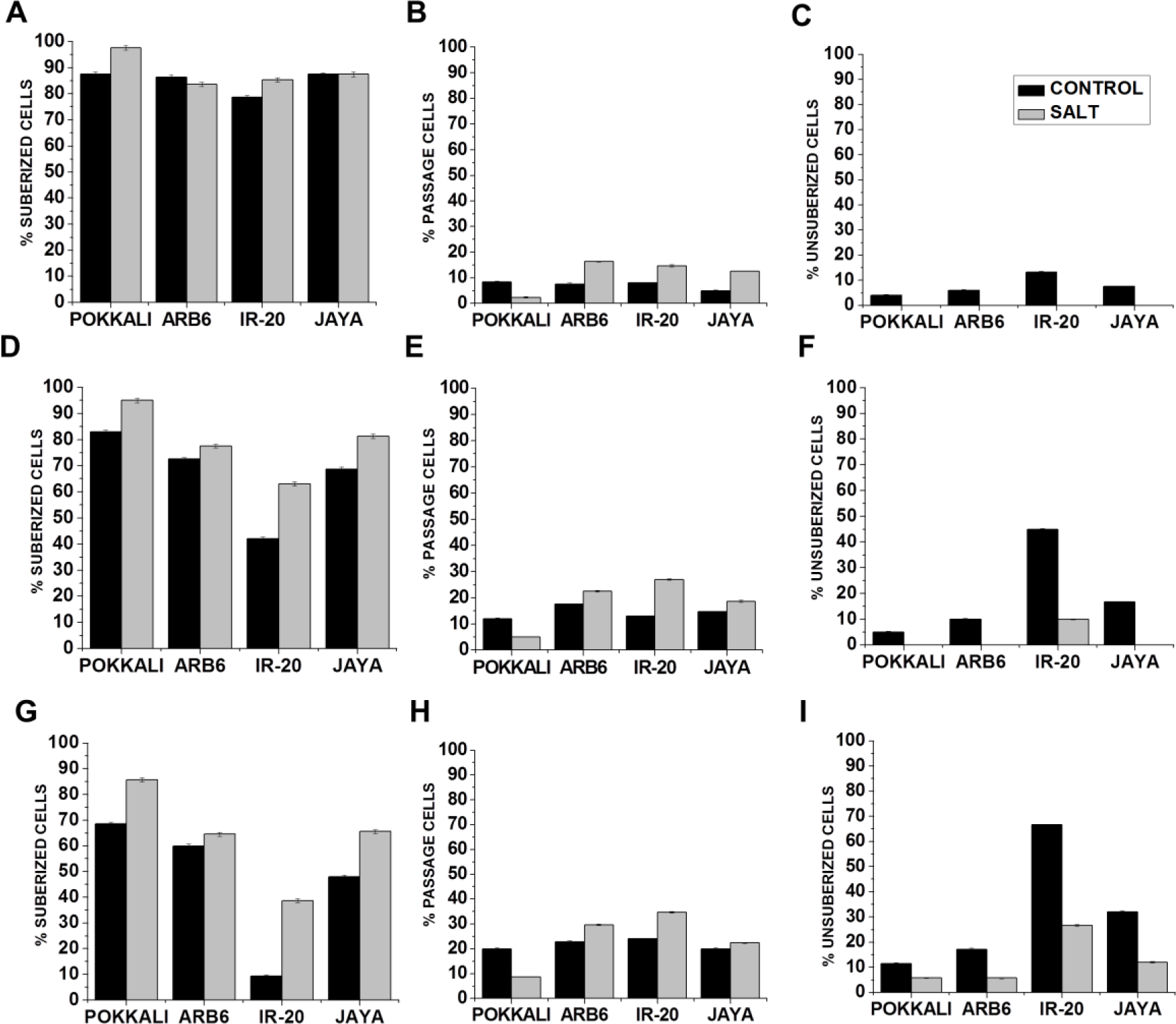
Quantification of suberization patterns in the endodermis to show suberized cells (having suberization on both sides and tangential areas), passage cells (with no tangential suberization) and unsuberized cells (without any suberin deposition) at (A - C) Base (D - F) Middle and (G - I) Tip of rice roots grown in control (well-watered) and salt (150mM NaCI) condition. Data represents mean (±SE; n = 6)

#### Na^+^ content in rice varieties under salt stress

The Na^+^ content was much higher under saline stress than in control samples in all four varieties, but was highest in IR-20 followed by Jaya and ARB6 (Figure 9A). It was least in Pokkali. Despite maintaining low Na^+^ content in the xylem sap, Pokkali had somewhat higher total shoot Na^+^ levels than the other three, which were comparable (Figure 9B). Apoplastic Na^+^, which we have previously shown to correlate well with survival under salt stress [27], showed a different trend. Pokkali had by far the lowest apoplastic Na^+^ and IR-20 had the highest Na^+^ content with ARB6 and Jaya being intermediate (Figure 9C). Much of the shoot Na^+^ in Pokkali appears to be present in the intracellular compartment with a negligible amount being sequestered in this compartment by IR-20 (Figure 9D). Our earlier studies would suggest that the bulk of the intracellular Na^+^ in Pokkali is sequestered in vacuoles [30]. Intracellular K^+^ levels were well controlled in all the four varieties under salt stress (Supplementary Information 8)

**Figure 9.**
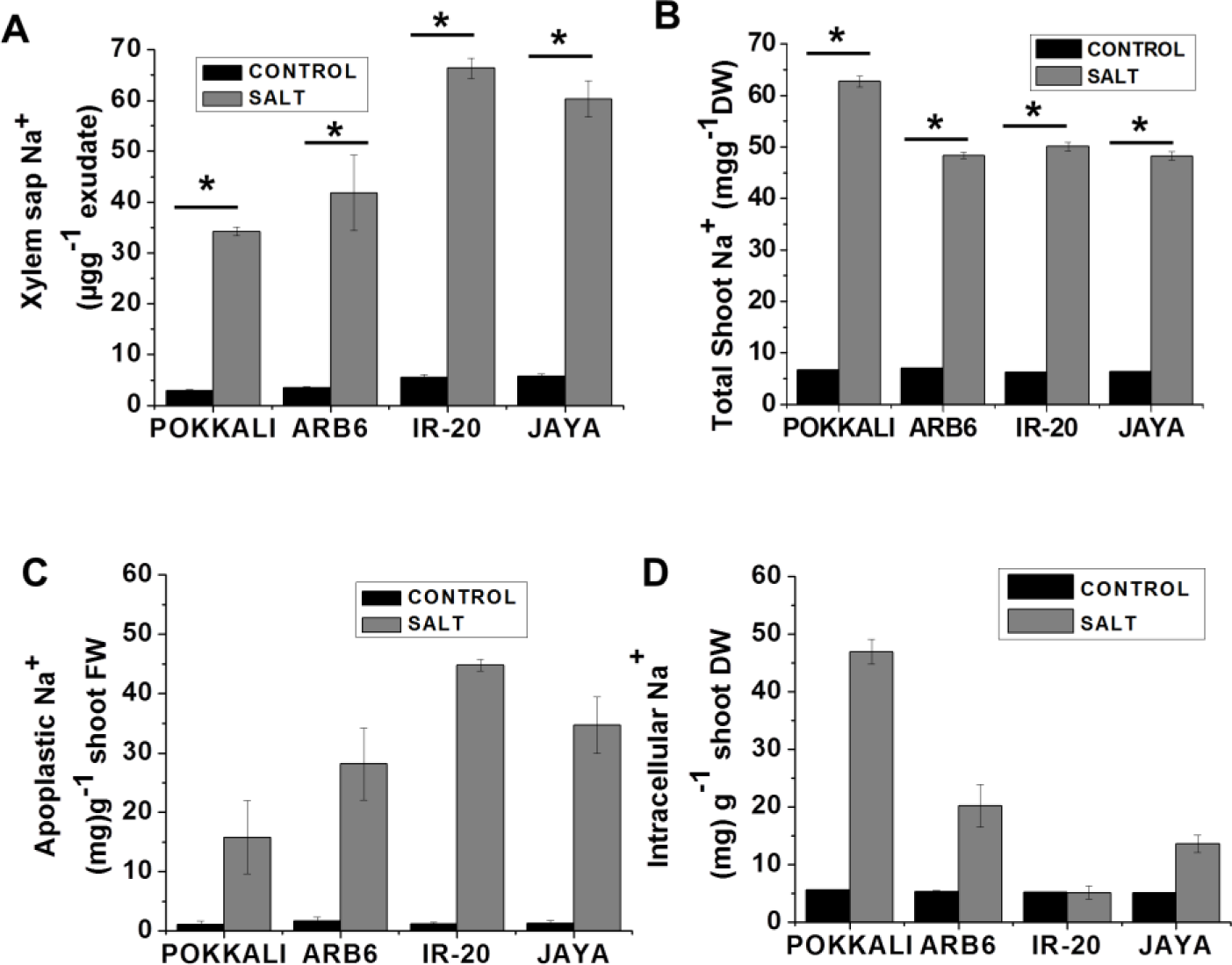
Na^+^ uptake by rice varieties. Plants were grown in PVC pipes for 38 days and then subjected to control (well-watered) or salt (150mM) for a week. (A) Na^+^ content in xylem sap. (B) Shoot total Na^+^ content. (C) Na^+^ content of shoot apoplastic fluid. (D) Intracellular Na^+^ content. After stress small fragments of shoots were rocked in double distilled water and supernatant was analyzed for apoplastic Na^+^ content. Simultaneously, shoots devoid of apoplastic Na^+^ were dried, powdered and suspended in distilled water. Na^+^ levels were estimated by flame photometry. Data represents mean (±SE; n = 6), Asterisk indicates differences between control and stressed at P < 0.05.

### Root hydraulic conductivity

Hydraulic conductivity decreased significantly under saline stress in all varieties except IR-20 (Table 1). Osmotically driven conductivity (*L*_0_) dropped by a factor of 2 or more in the cases of Pokkali, ARB6 and Jaya, ending at values of around 2 (ms^−1^) (Table 1). The decline was maximum for Pokkali (67 %) and less for ARB6 (60 %) and Jaya (50%). IR-20 finished the salt stress period with an osmotically driven hydraulic conductivity that was 65% greater than that of Pokkali. Similar declines were observed for external pressure driven hydraulic conductivity (*L*_pr_) with declines exceeding 50% for Pokkali and ARB6 while Jaya exhibited a decline of 40%. These three varieties had final conductivity values around 20 x 10^−8^m MPa^−1^ s^−1^. The decline in IR-20 was marginal.

### Expression of suberin synthesis genes in roots under drought and salt stress condition

To check the expression of suberin synthesis genes, total RNA from roots of control and stressed four rice varieties was analyzed by RT-PCR. Figure 10 shows expression of three *Elongases* (Os03g12030, Os02g11070 and Os06g39750) and one *cytochrome P450* (Os01g63540) after one week of drought and salt stress. *Elongase* (Os03g12030) was found to express equally in all four rice varieties under control, drought and salt stress. The expression of *Elongase* (Os02g11070) had dramatically reduced in all four varieties under drought stress compared to salt and control condition. While *Elongase* (Os06g39750) was found to express in all four varieties under salt stress, there was a dramatic decline in expression in Pokkali and ARB6 under drought stress. The expression of *cytochrome P450* (Os01g63540) was essentially the same in all four varieties under both drought and salt stress.

**Figure 10.**
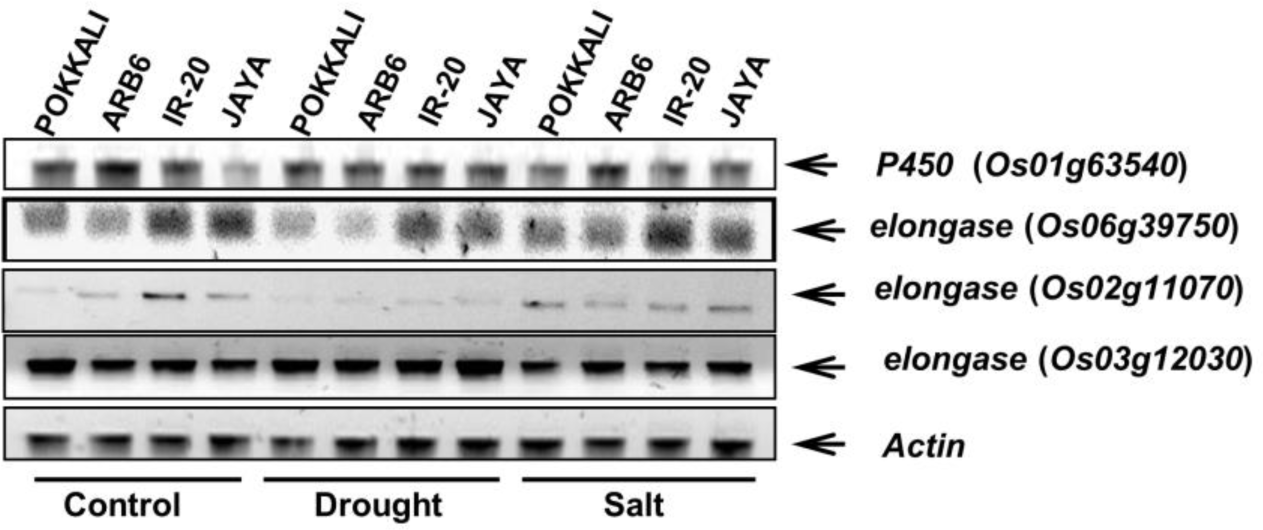
Expression of elongases and P450 in roots of four rice varieties after one week of drought and salt stress monitored by RT-PCR.

## Discussion

Salinity and drought are among the major factors limiting crop productivity worldwide. Both are essentially water deficit problems, although salinity has the added feature of Na^+^ toxicity. While some responses are common across several abiotic stresses, others are stress specific and may well prove counterproductive in dealing with others. It was to explore this possibility that we have subjected ―specialist‖ varieties reported to be tolerant to either drought or salinity to both stresses and contrasted their responses. We have used conditions where all the plants survived the stress protocol. Measurement of soil moisture content and electrical conductivity indicated similar and significant amounts of stress being experienced by all varieties in each protocol (Supplementary Information 1A, B, C, D, E). Photosynthetic rates declined significantly over the period of stress in all cases (Figure 1C and Figure 6C), the extent of decline being moderate for the tolerant varieties and large for the sensitive varieties. It was very surprising to find that both Pokkali and ARB6 maintained good photosynthetic rates when subjected to the stress they had not been selected for.

Root architecture and physiology may be expected to play key roles in surviving drought and salinity. Indeed long and prolific root systems are traits selected for while breeding for drought tolerance[14,46,47]. Minimization of apoplastic flow that bypasses hydrophobic barriers in the endodermis is a valuable trait in selecting for salt tolerant lines [48,49]. This trait could have deleterious effects on root hydraulics and may not be conducive to survival under drought. Thus varieties chosen for their stellar performance in one stress regime may not necessarily do well under a contrasting stress. Features such as the activity and localization of ion transporters or aquaporins may be critical in one regime while playing minor roles in the other. Our current study has focused on features affecting fluid uptake and delivery.

Pokkali is known to develop extensive hydrophobic barriers around both the endodermis and the exodermis constitutively. The unexpected finding that Pokkali does well under drought stress prompted the question as to whether the two successful varieties deployed different strategies, one enhancing water use efficiency while the other resorts to water mining. The observation of a large differential between canopy and ambient temperature in both ARB6 and Pokkali under drought stress strongly suggests evaporative cooling and hence a substantial transpirational stream even in Pokkali (Figure 2B), which is consistent with water mining. High rates of photosynthesis and xylem fluid exudation by both varieties under drought (Fig. 1c and Figure 2C) support this hypothesis. Vigorous transpiration in the face of drought requires access to subsurface water. Both Pokkali and ARB6 have long roots that elongate further under drought stress ending up over 50% longer than IR-20 roots (Figure 1G), facilitating access to deeper layers.

Uptake is driven by accumulation of osmolytes in both ARB6 and Pokkali. Accumulation of osmolytes is much higher in the face of drought stress (Figure 2D) than in salinity in case of ARB6 (Figure 7C), while the opposite is true for Pokkali. Jaya, which was earlier shown to adapt well to salinity, also accumulates a modest amount of osmolytes under saline stress, but not under drought. IR-20, which also adapted to saline stress by building hydrophobic barriers and restricting bypass flow under hydroponic culture [23], did not accumulate any osmolytes under either drought or salinity stress.

The water mining strategy is likely to result in roots growing deep into hypoxic environments. Transporting oxygen down from the shoots requires extensive aerenchyma formation, while suberization of the exodermis would minimize radial oxygen loss to the surrounding soil. Extension of the size and extent of aerenchyma is seen in both ARB6 and Pokkali in response to stress (Supplementary Information 4), along with substantial suberization of the exodermis (Supplementary Information 6). This adaptation would serve both to reduce the number of cells drawing on scarce resources under stress as well as supplying oxygen to distal portions of the root.

The hydraulic conductivity we report is that measured with roots immersed in standard growth medium in all cases. The osmotically-driven conductivity (*L*_0_) is measured in the absence of any external pressure and reflects the hydraulics of the symplastic cell-to-cell pathway which is strongly influenced by aquaporin conductivity [50]. In addition, fluid uptake may be expected to reflect differences in water potential across the plasma membrane. The large increase in osmolyte content under drought would lead to an expectation of enhanced driving force for water with the roots in growth medium and hence enhanced *L*_0_. The underwhelming increase in *L*_0_ is puzzling. Sap exudation rate, on the other hand, is measured with plants in soil and reflects a combination of *L*_0_ and the difference of water potential between root cells and the soil surrounding the root. The fact that ARB6 and Pokkali maintain moderate rates of xylem sap exudation even in drought or salinity must then be ascribed to the osmotic adjustment made in the two cases, together with the reduction in completely suberized cells in these two varieties.

Pressure-driven hydraulic conductivity *L*_pr_ reflects the hydraulics of the apoplastic pathway, primarily due to suberization patterns of the endodermis [51,52]. Exodermal suberization has been shown not to influence *L*_pr_ significantly in maize [53] and rice [54]. Small, but significant increases in *L*_pr_ in ARB6, Pokkali and Jaya contribute to enhanced fluid flow through the apoplastic pathway under drought (Table 1). This corresponds well with the decrease on overall suberization and increase in passage cell number under drought in these varieties (Figure 5 and Supplementary Information 6). Conversely, the lack of such change in suberization patterns in IR-20 is echoed in the finding that *L*_pr_ is also unaffected under drought (Table 1). Salinity induces a substantial decrease in osmotic *L*_0_ suggesting either a decrease in aquaporin expression or activity or both. A concomitant decrease in hydrostatic *L*_pr_ correlates well with the increase in completely suberized cells at the cost of unsuberized cells in the endodermis of Pokkali, the most successful variety under saline stress (Figure 8). ARB6 and Jaya also increase suberization of the endodermis but to a smaller extent. Again, maintenance of xylem sap exudation despite a reduction in *L*_0_ is strongly correlated with osmolyte accumulation in both Pokkali and ARB6.

It is worth noting differences between the behavior of the plants grown in soil and hydroponics. Roots of all varieties are extensively suberized when grown in well-watered soil, as opposed to hydroponics [23]. Further, the barriers formed in response to a week of saline stress do not completely eliminate osmotically driven xylem fluid exudation in soil (Figure 7B) as it does in hydroponics [23]. A week of conditioning stress under hydroponic culture rendered IR-20 almost as tolerant to salt as Pokkali, whereas the extent of adaptation when grown in soil appears to be less effective. It would thus appear that not only are modest barriers already in place under control conditions when grown in soil, but that the extent to which they are fortified in response to salinity stress is much less than under hydroponics. Extension of results from hydroponics to field conditions may need to be taken with a pinch of salt.

The impressive rates of exudation seen in ARB6 and Pokkali, especially under drought, could be attributed to the presence of passage cells. Indeed, the dramatic increase in passage cells is one of the most striking features of the drought tolerance response of ARB6. It may be noted that while passage cell numbers increase under drought stress, cells with unsuberized radial walls are completely eliminated in Pokkali and largely so in ARB6 under salt stress. Thus the pattern of suberization is dramatically different in the face of different stresses. This is illustrated in Figure 11, which shows a near linear variation of xylem sap exudation rates and xylem sap Na^+^ content with the number of passage cells under drought and salinity (Figure 11A, B). The only variety to do well under salt stress is Pokkali, which completely eliminates unsuberized cells in the base and middle regions of the endodermis and also significantly reduces passage cells. ARB6 also eliminates unsuberized cells, but slightly increases passage cells in this condition. The dramatic reduction in the number of completely suberized cells in the base region of both endodermis and exodermis of ARB6 and Pokkali in response to drought is puzzling, as it implies the removal of suberin deposits. No mechanism for depolymerization of suberin has thus far been reported. This may well prove to be a suitable system to explore such mechanisms. These findings demonstrate that suberin deposits are laid down in very different patterns in response to the two stresses. Intriguingly, both ARB6 and Pokkali are capable of remodeling their hydrophobic barriers appropriately in both cases while IR-20 is not.

**Figure 11.**
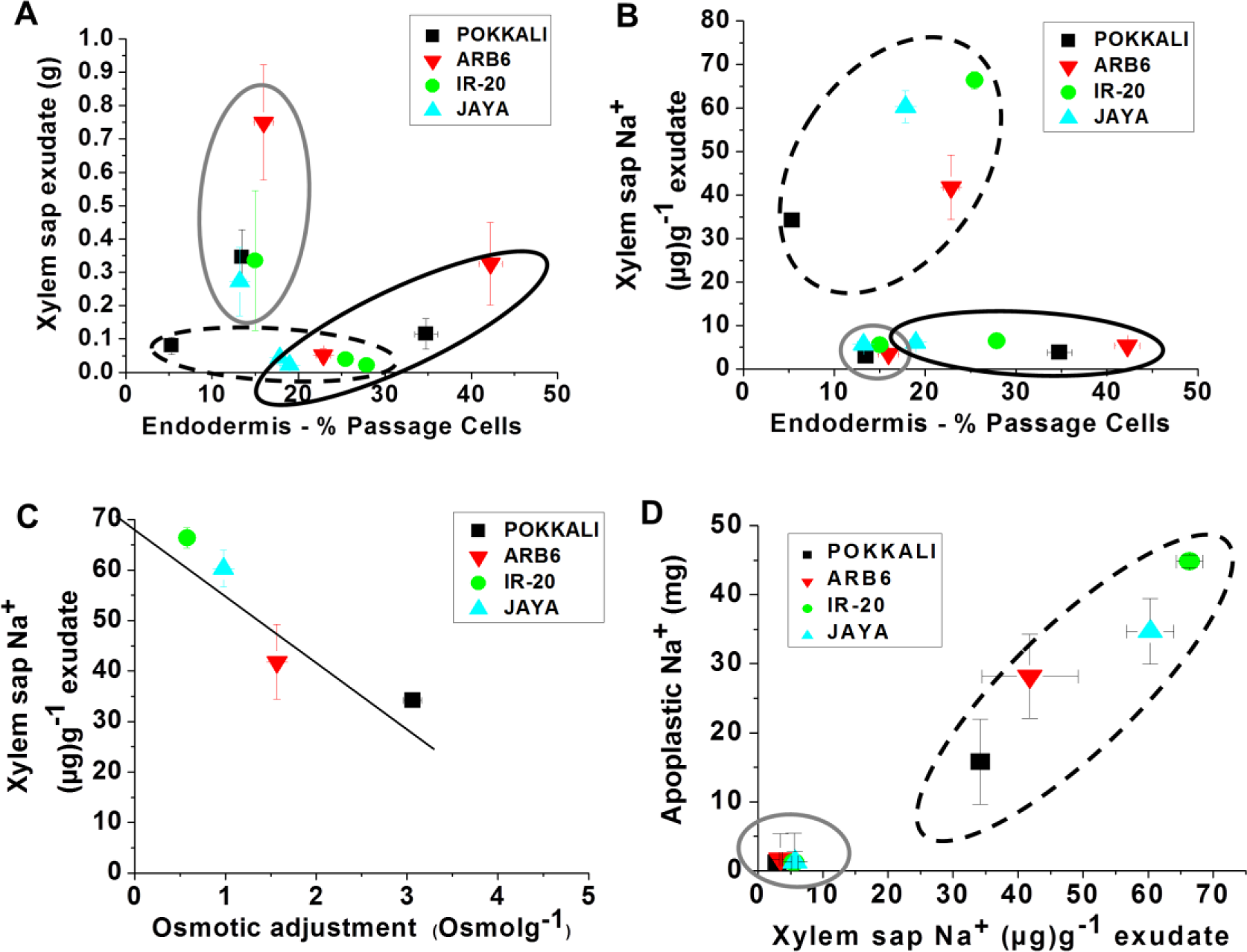
(A) Relation between passage cells of endodermis and xylem sap uptake. (B) Relation between passage cells of endodermis and xylem sap Na^+^ content. (C) Relation between xylem sap Na^+^ content and osmotic adjustment. (D) Relation between xylem sap Na^+^ content and apoplastic Na^+^. Data represents mean (±SE; n = 6). (Grey circle - Control, Black circle - Drought and Dashed circle - Salt)

The elimination of unsuberized cells in the endodermis of Pokkali, ARB6 and Jaya under salt means that fluid entry over much of the length of the root occurs through passage cells. This ensures that solutes have to cross at least two sets of plasma membranes before entering the xylem stream. Solute movement across cell membranes should be subject to selectivity of the relevant transporters. The large build-up of osmolytes in Pokkali compensates for the restriction in fluid entry points. It also appears to contribute to minimizing Na^+^ loading of the xylem sap as we observe a nearly linear decline in the Na^+^ content of sap with osmolyte accumulation (Figure 11C). This reduction in xylem sap Na^+^ is in turn correlated with limiting Na^+^ in the shoot apoplast (Figure 11D), which is tightly correlated to survival [28]. A significant part of the total shoot Na^+^ in Pokkali is located in the intracellular fraction. Our earlier studies on cellular mechanisms of combating salinity would suggest that this reflects the sequestering the Na^+^ in vacuoles, thereby minimizing the amount present in the apoplastic fraction. This sequestration mechanism appears not to function effectively in IR-20.

In RT-PCR data, the enhanced mRNA expression levels of *P450* and *elongases* (39750, 11070 & 12030) under salinity stress in all four varieties (Figure 10) was consistent with the observations of [23,55]. Whereas, there was significantly low expression of *elongase* (11070) and *elongase* (39750) under drought stress (Figure 10). The suberization pattern and RT-PCR data appears to suggest that the suberin synthesis pathway and deposition in tolerant and sensitive rice varieties under salinity and drought stress could be different.

This study comparing the physiological responses of four cultivars varying widely in tolerance to drought and salinity clearly identifies several features of roots that contribute to survival under these stresses. An extensive root system facilitates water mining in drought. Aerenchyma formation reduces the number of resource consuming cells as well as providing a conduit for oxygen to distal portions of the root; suberization of the exodermis minimizes loss of the transported oxygen. The pattern of suberization differs greatly between drought and salinity, but in both cases passage cells facilitate fluid uptake. In the latter case, the only endodermal cells lacking tangential suberin are passage cells – thereby ensuring solute selectivity. Hydrostatic conductivity changes are prominent under salinity, restricting fluid flow to the symplastic pathway and where solutes can be sifted by selective transporters. Uptake is driven by osmolyte loading, which also contributes to reduction of Na^+^ uptake. Our study is purely correlative. However, the use of four varieties with a wide span of sensitivity to each stress provides significant spread in the stress– driven parameters monitored, thereby strengthening the conclusions drawn. We had hypothesized that specialist varieties would do well under ―their‖ respective stress, but would be sensitive to the other stress; and that the responses would be graded according to the degree of tolerance/sensitivity among the four cultivars studied. The latter expectation was indeed upheld with photosynthetic rates spanning a wide range at the end of each stress protocol. The former expectation was based on an assumption that the ―specialists‖ are restricted to one response to all stresses. This is clearly not the case.

In conclusion, the set of adaptations we describe appears to make a significant contribution to the ability of rice plants to survive water deficit conditions in both drought and salinity stress. The study has provided experimental evidence to validate the large quantity of empirical field data related to drought tolerance of ARB6 and salt tolerance of Pokkali. Testing the effectiveness of remodeling suberization patterns will require tools to perturb such patterns. Mechanisms for the detection of stress and the regulation of the responses deployed are areas for further work as is the search for additional mechanism underlying drought and stress tolerance at the root level.

## Acknowledgement

We are grateful to Dr. Shivaprasad and Dr. Kalika Prasad for critical reviews of this manuscript. We also thank Ashvini Kumar Dubey, Anirban Baral and Raveendra GM for their timely advice and help. The Central Imaging & Flow Cytometry Facility (CIFF) at National Centre for Biological Sciences (NCBS) Bangalore is gratefully acknowledged. The work was performed with internal NCBS funding. The funding from Department of Biotechnology (DBT), India is also highly acknowledged.

