## Supplemental Figures for "Fluid Uptake Pathways Are Differentially Modified in Rice (*Oryza sativa* L.) in Response to Drought and Salinity"

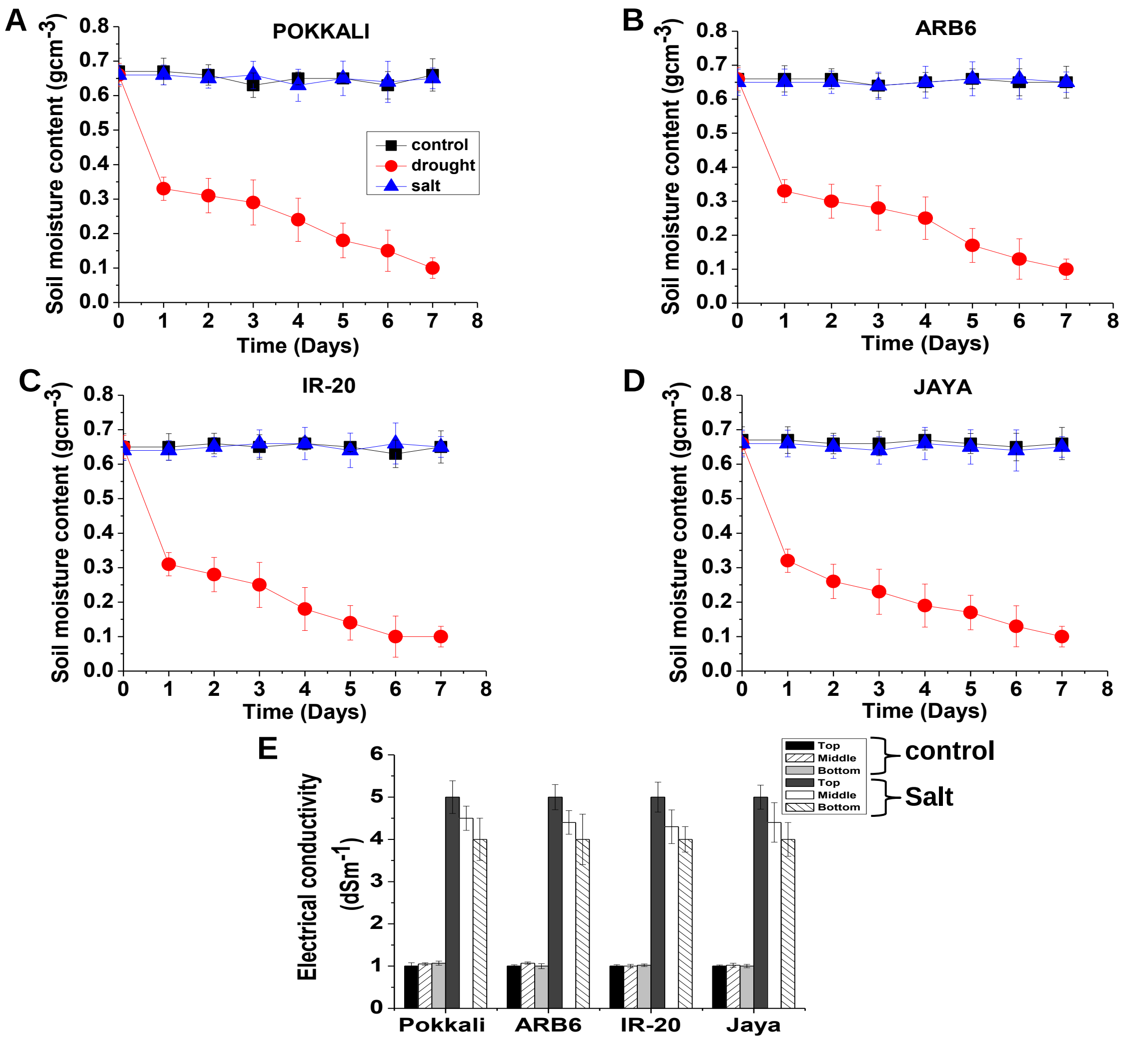


**Supplementary Information 1** Soil moisture content measured under control (well-watered), drought and Salt conditions in (A) Pokkali (B) ARB6 (C) IR-20 (D) Jaya during one week of stress period. (E) Electrical conductivity of soil under control and salt conditions in all four varieties. Soil in PVC pipes was varieties irrigated with salt water (150mM NaCl) followed by normal water irrigation to ensure the whole root system is exposed to salt stress. Electrical conductivity (EC) of soil samples collected from top, middle (Mid) and bottom (bot) portion of pipe was measured. 1g of soil was dissolved in 5ml of deionized double distilled water, supernatant was used to estimate EC.

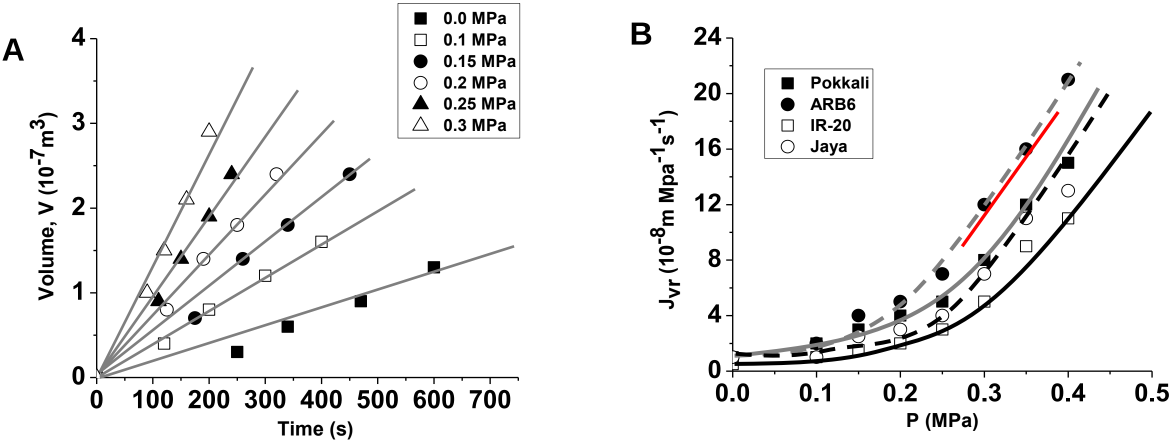


**Supplementary Information 2** Results from a typical steady-state experiment for measuring root hydraulic conductivity ($L_{pr}$). One week stress of drought (no irrigation) and salinity (150mM NaCl) was given to 38 day old plants of four rice varieties. Before starting the measurements, soil was washed off the roots and then transferred to nutrient solution. (A) Exuded xylem sap in Pokkali in the presence of hydrostatic pressure gradients ($P$) is plotted against time. (B) Steady-state water flow per unit surface area of the root system ($J_{vr}$) as a function of applied driving force. $L_{pr}$ values were calculated from slopes of the linear ranges (red line) of $J_{vr}$ ($P$) curves. Lines in A are fitted lines, after estimating the slopes of these lines the values of $J_{vr}$ were calculated. Lines passing through the data points in B are intended to guide the eye.

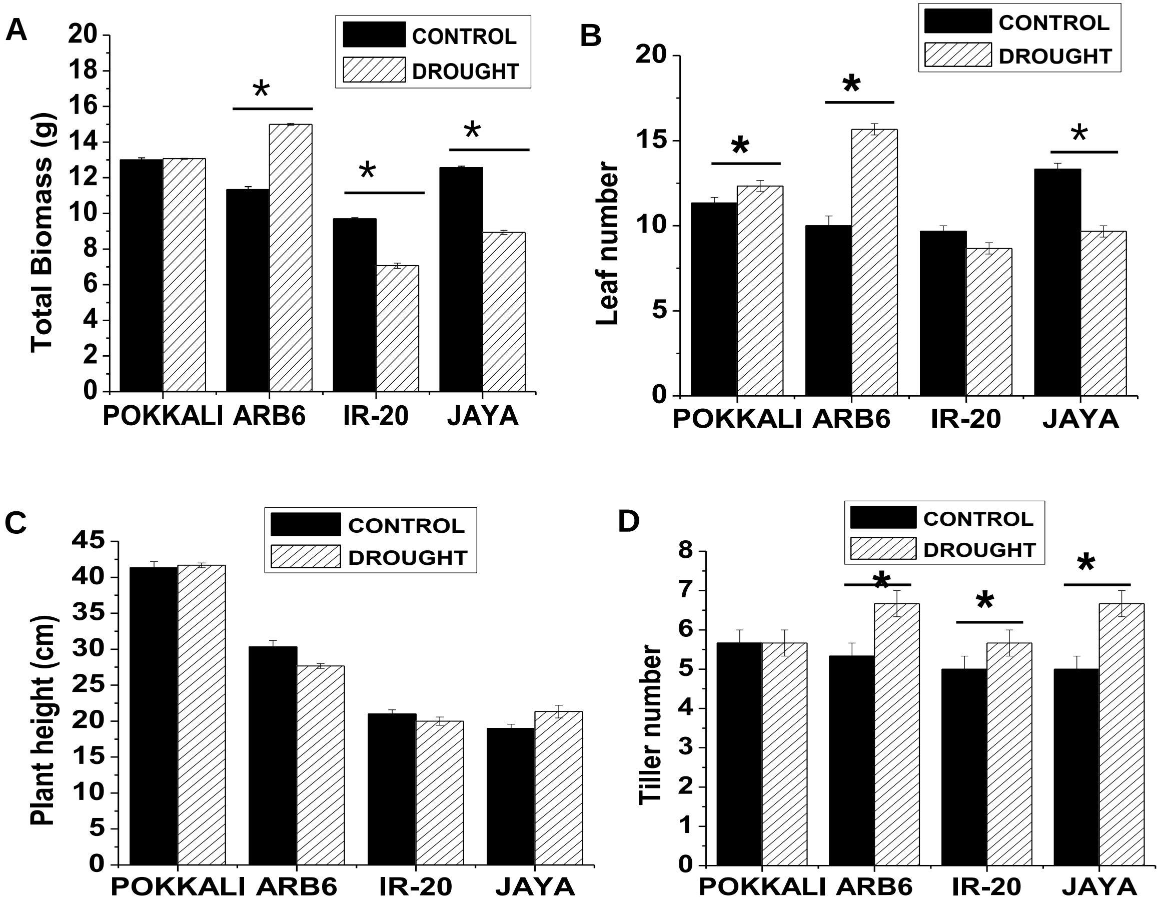


**Supplementary Information 3** Plants were grown in PVC pipes for 38 days and subjected to drought (no irrigation) condition for a week. (A) Total biomass (B) Leaf number (C) plant height and (D) tiller number were measured. Data represents mean (±SE; n = 6), Asterisks indicate differences between control and stressed at *P < 0.05 and **P < 0.01.

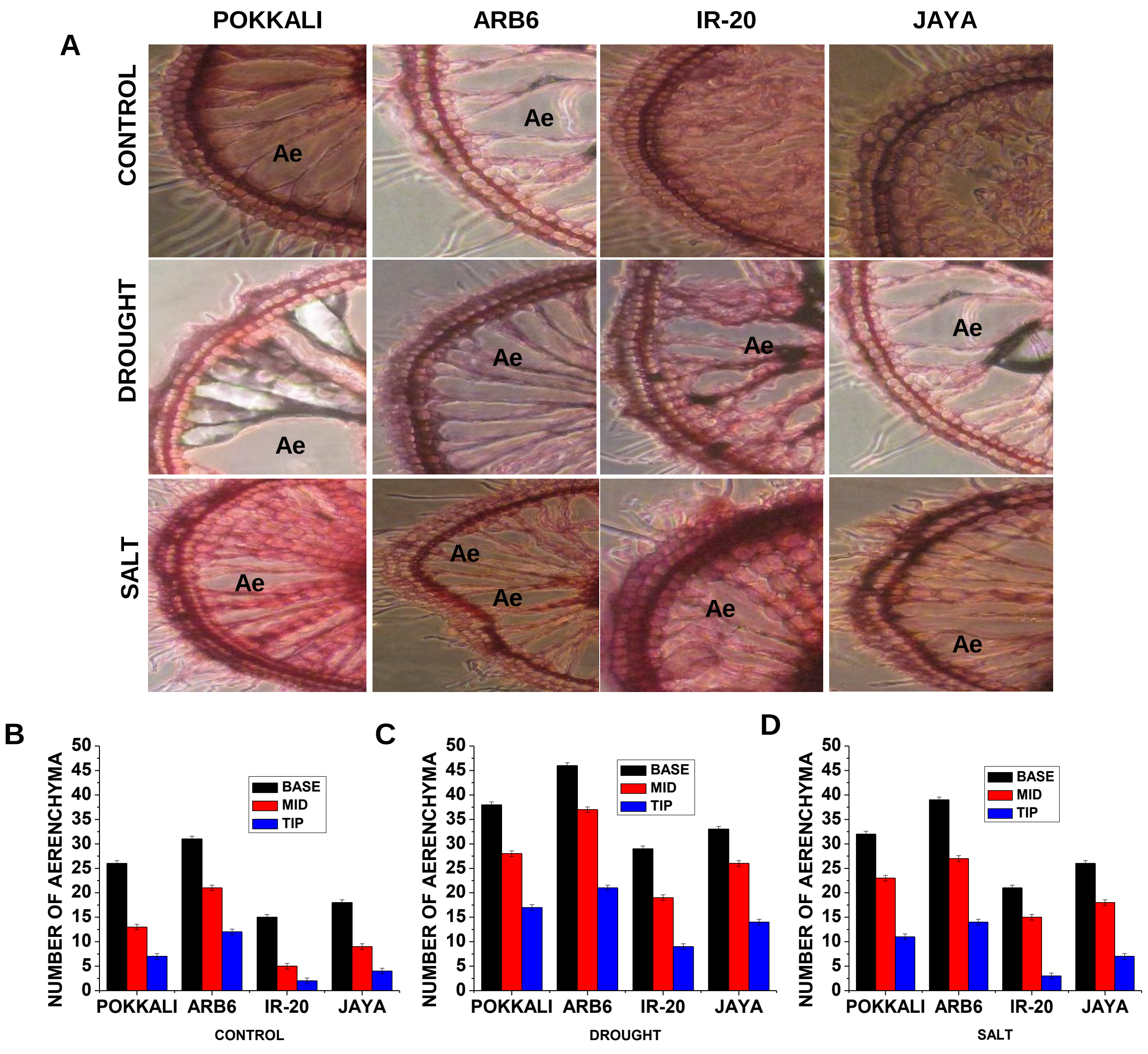


**Supplementary Information 4** (A) Aerenchyma (Ae) images in roots of Pokkali, ARB6, IR-20 and Jaya. Quantification of number of aerenchyma in base, middle and tip region of these plants grown in (B) Control (well-watered) (C) drought and (D) salt (150mM) condition.

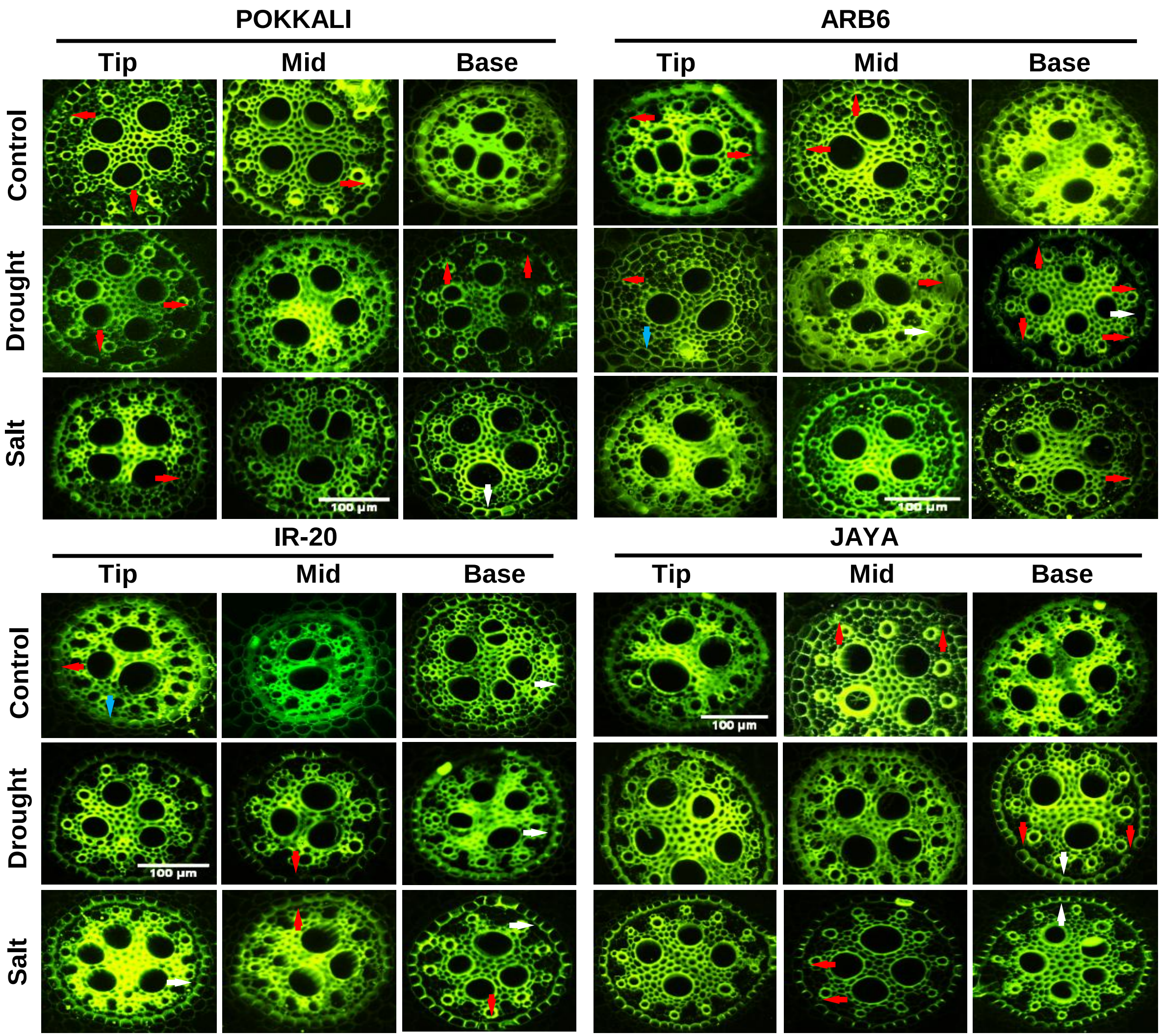


**Supplementary Information 5** Images of root sections showing passage cells and suberin deposits (Casparian strip) in root endodermis of four rice varieties grown in soil filled PVC pipes for 45 days and subjected to control (well-watered), drought (no irrigation) and salt stress (150mM) conditions on 38$^{th}$ day for a week. Roots were washed, divided into three zones (Tip, Mid and Base) and cut into thin sections (200µm), then stained with berberine-aniline blue and imaged using Olympus FV1000 confocal microscope, 488nm laser was used for excitation. Red, white and blue arrows show passage cells (cells with no tangential suberin deposition), suberin deposits and unsuberized cells respectively. *Bars* 100µm

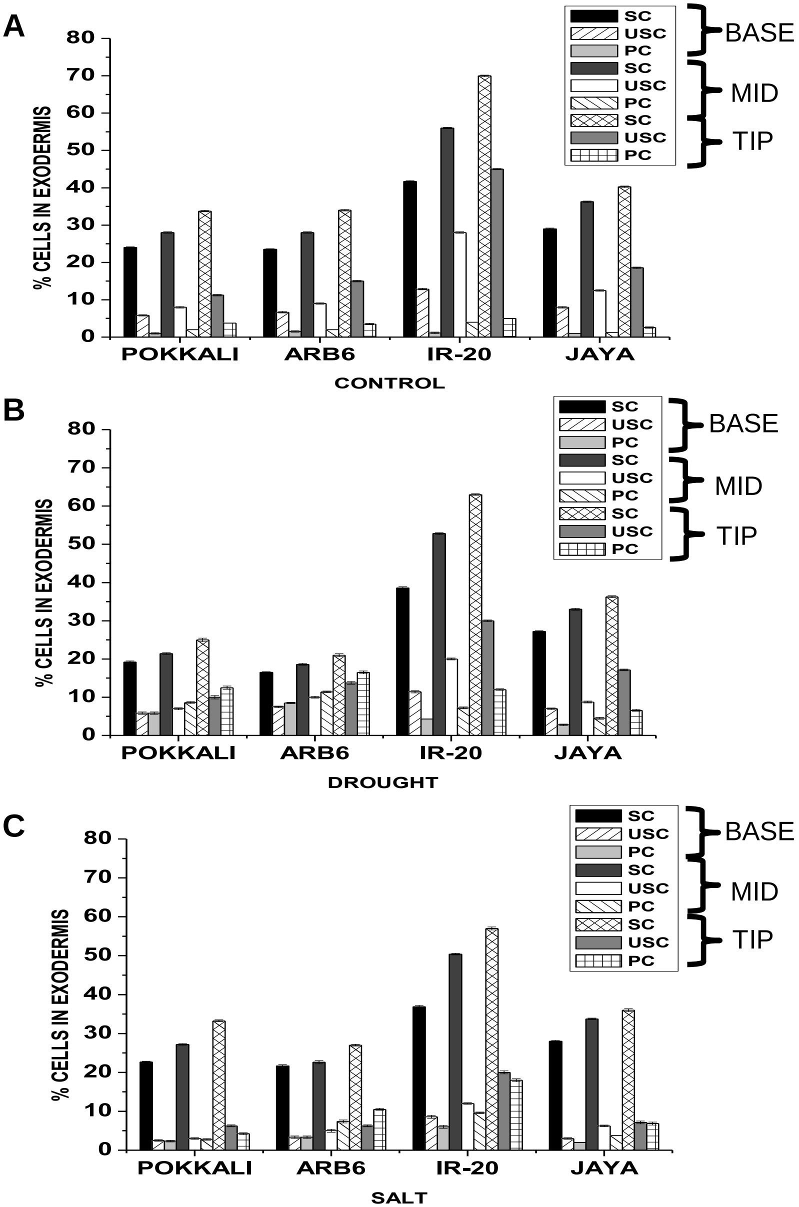


**Supplementary Information 6** Percentage of suberized cells (SC), unsuberized cells (USC) and passage cells (PC) in exodermis at base, middle and tip region of roots of Pokkali, ARB6**,** IR-20 and Jaya grown in (A) well-watered, (B) drought and (C) salt (150mM) condition.

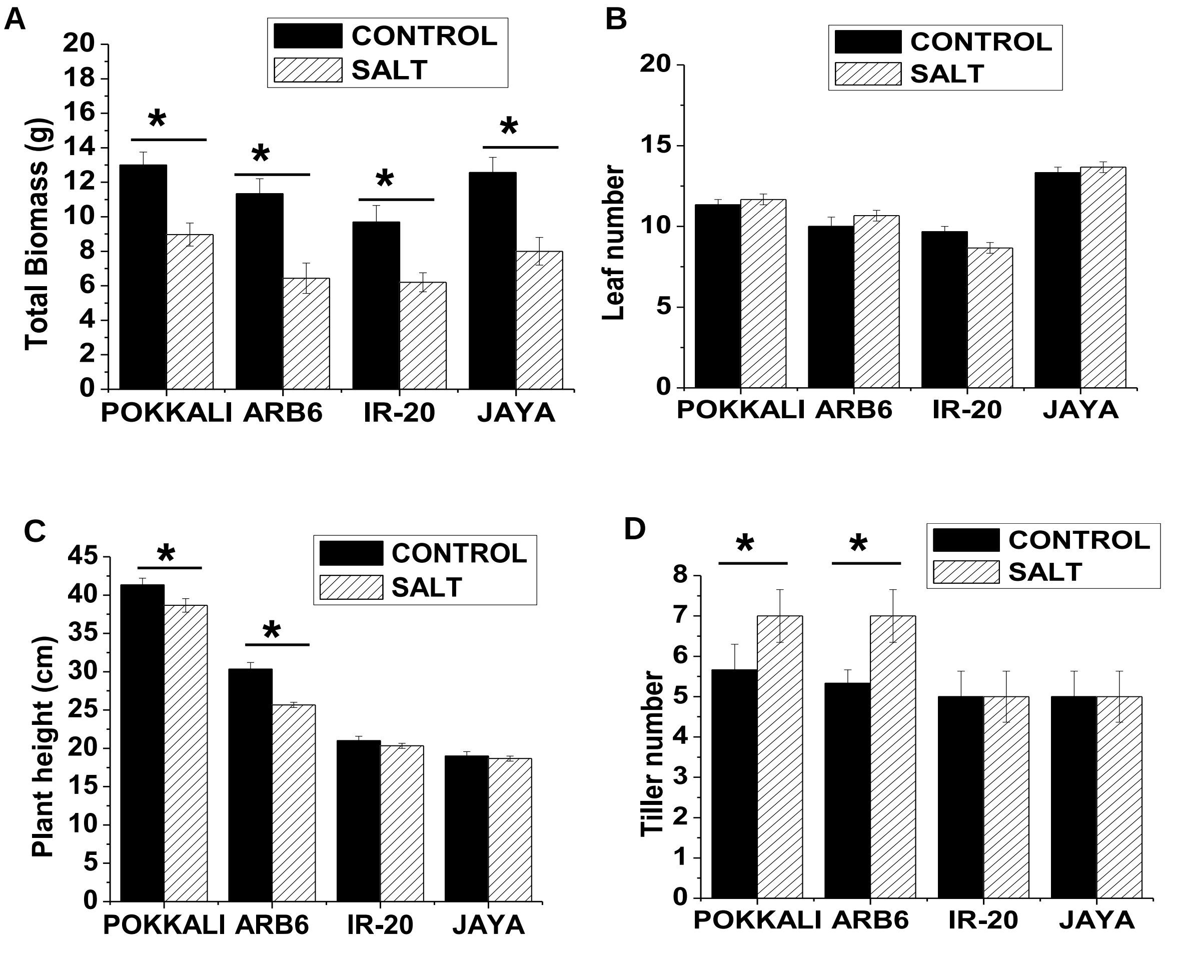


**Supplementary Information 7** Plants were grown in PVC pipes for 38 days and subjected to salinity (150mM NaCl) condition for a week. (A) Total biomass (B) Leaf number (C) plant height and (D) tiller number were measured. Data represents mean (±SE; n = 6), Asterisks indicate differences between control and stressed at *P < 0.05 and **P < 0.01.

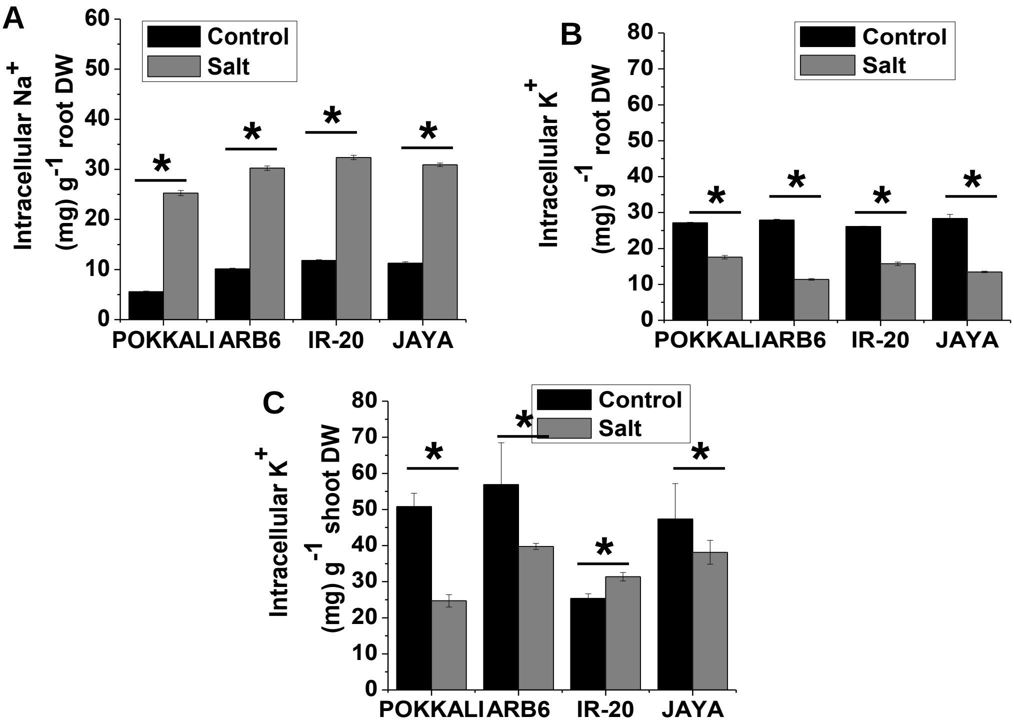


**Supplementary Information 8** $Na^+$ and $K^+$ uptake by rice varieties. Rice plants grown in PVC pipes for 45 days and subjected to control (well-watered) and salt stress (150mM) conditions for a week. (A) Intracellular $Na^+$ ions in root and (B,C) Intracellular $K^+$ ions in root and shoot respectively. After stress shoots were washed and cut into small fragments, then rocked in double distilled water for an hour and supernatant was analyzed for apoplastic $Na^+$ content. Simultaneously, shoots and roots devoid of apoplastic $Na^+$ were dried, powdered and digested in distilled water to estimate Intracellular $Na^+$ and $K^+$ levels by flame photometry. Data represents the mean (±SE; n = 6) with Asterisks indicating significant difference (*$P < 0.05$).
